# PAMPA: a software for peptide markers and taxonomic identification for ZooMS samples in Archaeology and Paleontology

**DOI:** 10.1101/2025.03.26.645441

**Authors:** Fabrice Bray, Hélène Touzet

## Abstract

ZooMS (Zooarchaeology by Mass Spectrometry) offers a rapid and cost-effective method for species identification of animal remains through peptide mass fingerprinting. After mass spectrum generation, a mostly used way to perform taxonomic identification is to compare the mass fingerprints to a reference database of diagnostic peptide markers to determine the species of origin. This analytical stage, however, is tedious and error-prone, often necessitating manual examination of spectra. In this paper, we present a comprehensive approach to automate and standardize the usage of peptide markers and the classification of ZooMS spectra. We have developed a software called PAMPA (Protein Analysis by Mass Spectrometry for Ancient Species), for which we demonstrate the effectiveness using a variety of spectral data from bone samples generated by MALDI-TOF and MALDI-FTICR. PAMPA is open-source and comes with a database of peptide markers and collection of curated COL1A1 and COL1A2 sequences. We believe it will be a valuable resource for the scientific community.

## 1. Introduction

In recent years, the interdisciplinary field of paleoproteomics has emerged as a powerful approach for analyzing animal remains and determining their taxonomic origins through the study of ancient proteins. Currently, the most widely used method in paleoproteomics is (ZooMS). ^1, 2^, that has been successfully applied to a diverse array of materials, including bone, teeth, hair, ivory, parchment, leather, and other soft tissues ^1, 3–7^ and animals, including mammals, reptiles, amphibians, birds, marsupials, and fishes ^8–25^ The first steps of the ZooMS process involves demineralization, followed by extraction and digestion, most often using the enzyme trypsin. The resulting peptides are analyzed by mass spectrometry, typically using a Matrix-Assisted Laser Desorption Ionization (MALDI) source coupled with a Time-of-Flight (TOF) detector, or more recently with a (Fourier-Transform Ion Cyclotron Resonance (FTICR) cell to obtain a MS profile.^26, 27^ The obtained mass profile predominantly contains type I collagen peptides, along with peptides from other proteins such as biglycan, alpha glycoproteins, keratins, and dermicins.^27^ Once the MS spectra have been generated, they need to be analyzed to identify the taxonomic origin of the sample. However, many collagen type I peptides are shared among closely related species, making species differentiation challenging, and software tools that are usually used in proteomics, such as the widely used search engine MASCOT, do not adapt easily to this task. In practice, in the vast majority of publications, the taxonomic assignment work is still done manually, focusing on identifying a selection of key diagnostic peptides rather. Such peptide markers have been empirically demonstrated to be taxonomically informative and thus serve as barcodes for the identification of particular taxa or species. One well-known set of peptide markers, extensively employed in the literature, is the collection of 12 peptides from COL1A1 and COL1A2 (Collagen 1 alpha 1, alpha 2) proposed by Buckley for mammalian species.^1, 28^ The drawback of manual assignment is that it is time-consuming, and raises issues of reliability, scalability and reproducibility. Efforts have therefore been recently made to propose bioinformatics approaches for automating this task. Bacollite introduced an algorithm for classifying a MALDI-TOF sample using the full proteomics profile of collagen type I sequences of each candidate species, provided that these sequences are available.^29^ SpecieScan used the principle of diagnostic peptides to automatically process MALDI-TOF spectra.^30^ In this article, we take the results of these works a step further and propose a comprehensive and user-friendly toolbox designed to efficiently handle a variety of tasks associated with ZooMS data and peptide markers. This toolbox, called PAMPA (Protein Analysis by Mass Spectrometry for Ancient Species), is based upon the concept of peptide markers, such as SpecieScan, and also allows the use of the full proteomics profile such as Bacollite. In both cases, it enables rapid and straightforward species identification from MALDI mass spectra. Additionally, PAMPA allows users to employ their own sets of peptide markers and offers functionalities to generate customized markers either by sequence similarity from existing markers, or from literature data in an assisted manner. We demonstrate its applicability on a series of case studies, and have used it to automatically build a large collection of peptide markers for mammals. We truly believe that PAMPA can help ensure consistency and reliability in ZooMS analyses, while allowing us to scale with a large number of samples.

## 2. Results

PAMPA is built around the central concept of *peptide tables*, that are used to compile peptide markers and their masses.

### 2.1 Peptide tables

In essence, a peptide table is a database of peptides described by either their sequence or mass, or both, and coupled with taxonomic information. It also contains optional metadata that eases readability and traceability: taxonomic rank, taxonomic identifiers, description of post-translational modifications applied to the peptide, marker name, peptide mass, gene name, position of the peptide within the helical region (following the nomenclature of Brown *et al*.^28^), sequence identifier(s) of the protein sequence from which the peptide is derived, start and end positions of the peptide within the protein sequence, status of the peptide regarding cleavage, and any additional comments about the marker. These peptide tables serve two purposes. First, they are an exchange format to describe peptide markers and their properties. Second, they can be passed as a parameter to the program to perform taxonomic assignment.

### 2.2 Creation of peptide tables

The construction of a peptide table can be achieved through various means with PAMPA. The first way is to start from data from the literature, and to build an initial peptide table by hand, containing partial information such as mass or marker sequence for a given species. PAMPA can then automatically complete missing fields. It can compute masses for peptides that lack this information. Moreover, if type I collagen sequences are provided, PAMPA can calculate peptide positions or, conversely, deduce the sequence and mass of a peptide marker from its positions in the sequence (Figure 1, A). It can also deduce the peptide sequences, together with their PTMs and their positions, from the mass. The second way to build a peptide table is to use a set of protein sequences as an entry point and to generate tryptic peptides from it. This scenario is useful when no marker peptide is reported for the species. This is achieved by performing systematic in-silico digestion and mass computation to construct a peptide table (Figure 1, B). The digestion allows up to one miss cleavage, and hydroxylation of prolines are automatically inferred. For that, the number of hydroxyprolines is determined empirically using the following formula: Let ‘p’ represent the total number of prolines in the peptide, and ‘pp’ represent the number of prolines involved in the pattern ‘GxP’. If the difference ‘p-pp’ is less than 3, then ‘pp’ hydroxyprolines are applied. If ‘p-pp’ is 3 or greater, ‘pp’ hydroxyprolines and ‘pp+1’ hydroxyprolines are applied. The set of tryptic peptides can then be filtered out by PAMPA in two ways. First, when mass spectra are available for the organism of interest, PAMPA allows to select only peptides whose mass matches with observed peaks. The list of tryptic peptides can also be refined by establishing homology with known peptide markers. In this case, the comparison is performed by sequence comparison allowing for mismatches. This approach is better than the usual use of BLAST, as it ensures that the cleavage sites are preserved and does not require any prior alignment between the sequences, meaning that it can successfully handle partial incomplete sequences or isoforms exhibiting large insertions or deletions. In all cases, peptide tables need to be constructed only once and can be edited manually. In particular, it is possible to modify peptide sequences, to authorize other PTMs as hydroxyproline, such as deamidation of asparagine and glutamine, or phosphorylation of serine, threonine, and tyrosine. The mass is then automatically updated.

**Figure 1:**
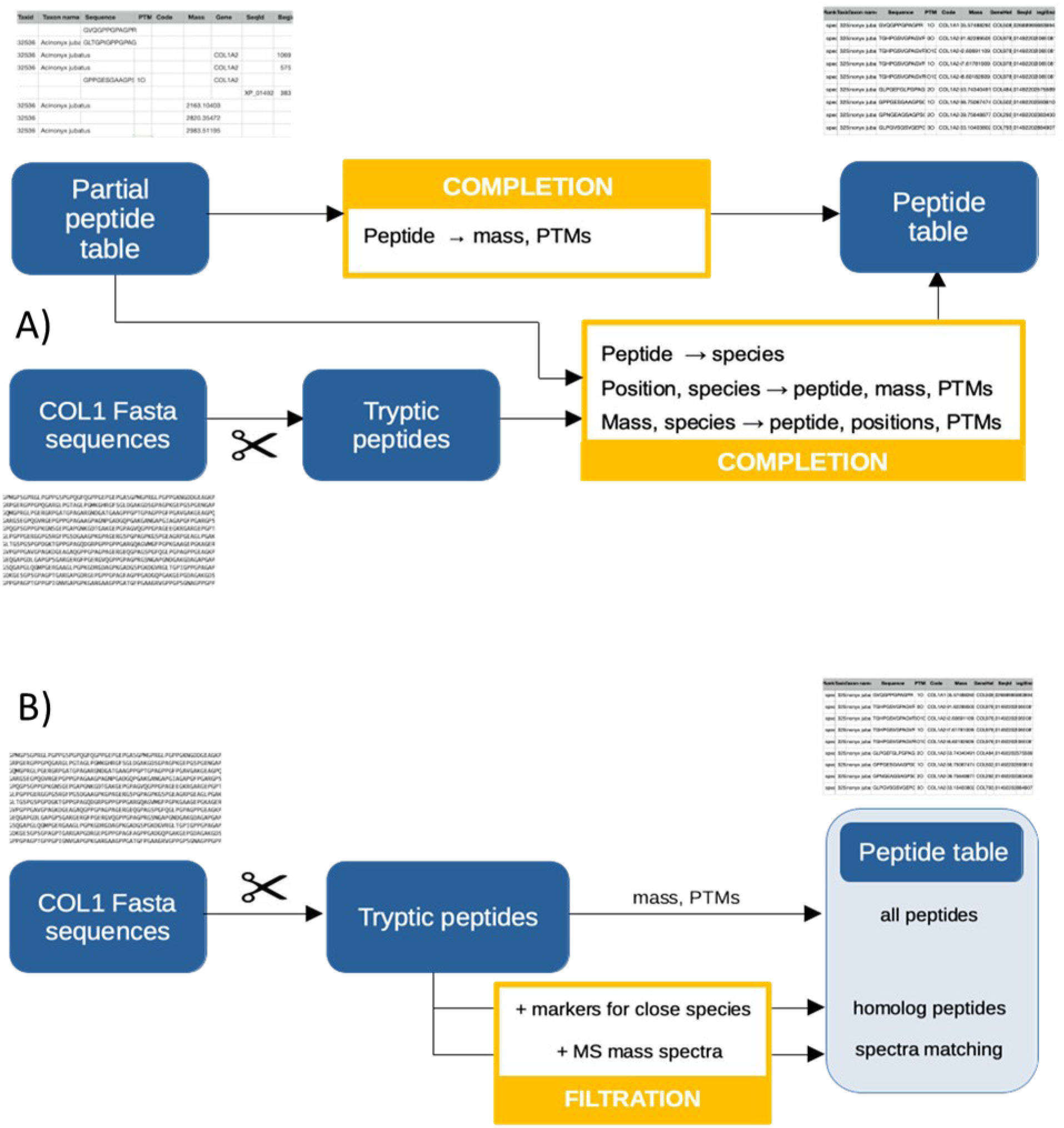
Two main ways to build peptide tables with PAMPA. A) by completion of a peptide table manually created from partial information found in the literature and automatically supplemented using Fasta sequences. B) by filtration from the full proteomic profile, either by homology or from model spectra.

### 2.3 A comprehensive peptide table for mammals

We used PAMPA to construct a reference peptide table that gathers ZooMS peptide markers for a large collection of mammal species: COL1A2 978-990, COL1A2 978-990 +16, COL1A2 484-498, COL1A2 502-519, COL1A2 793-816, COL1A1 586-618, COL1A1 586-618 +16, COL1A2 757-789, COL1A2 757-789 +16, COL1A1 508-519 and COL1A2 292-309 peptide markers (also named A, A’, B, C, D, F, F’, G, G’, P1, P2 respectively). We selected all species having both COL1A1 and COL1A2 sequences available in the NCBI database, and then further refined this selection to keep only sequences whose mature region is full-length. This makes a total of 418 COL1A1 and COL1A2 amino acids sequences coming from 209 species, whose taxonomic distribution is given in Table 1. Among these species, 55 have proteomics data available in the Googlesheet compilation “ZooMS Markers - Published data” maintained by S. Presslee (https://docs.google.com/spreadsheets/d/1ipm9fFFyha8IEzRO2F5zVXIk0ldwYiWgX5pGqETzBco/edit?gid=1005946405#gid=1005946405). In these cases, we have generated the corresponding peptide sequences with PAMPA from target masses, and compared them to the LC-MS/MS sequence when available. For the other species, we have inferred both sequences and masses by homology with PAMPA, automatically checking the correctness of the helical position and the tryptic cleavage.

**Table 1:**
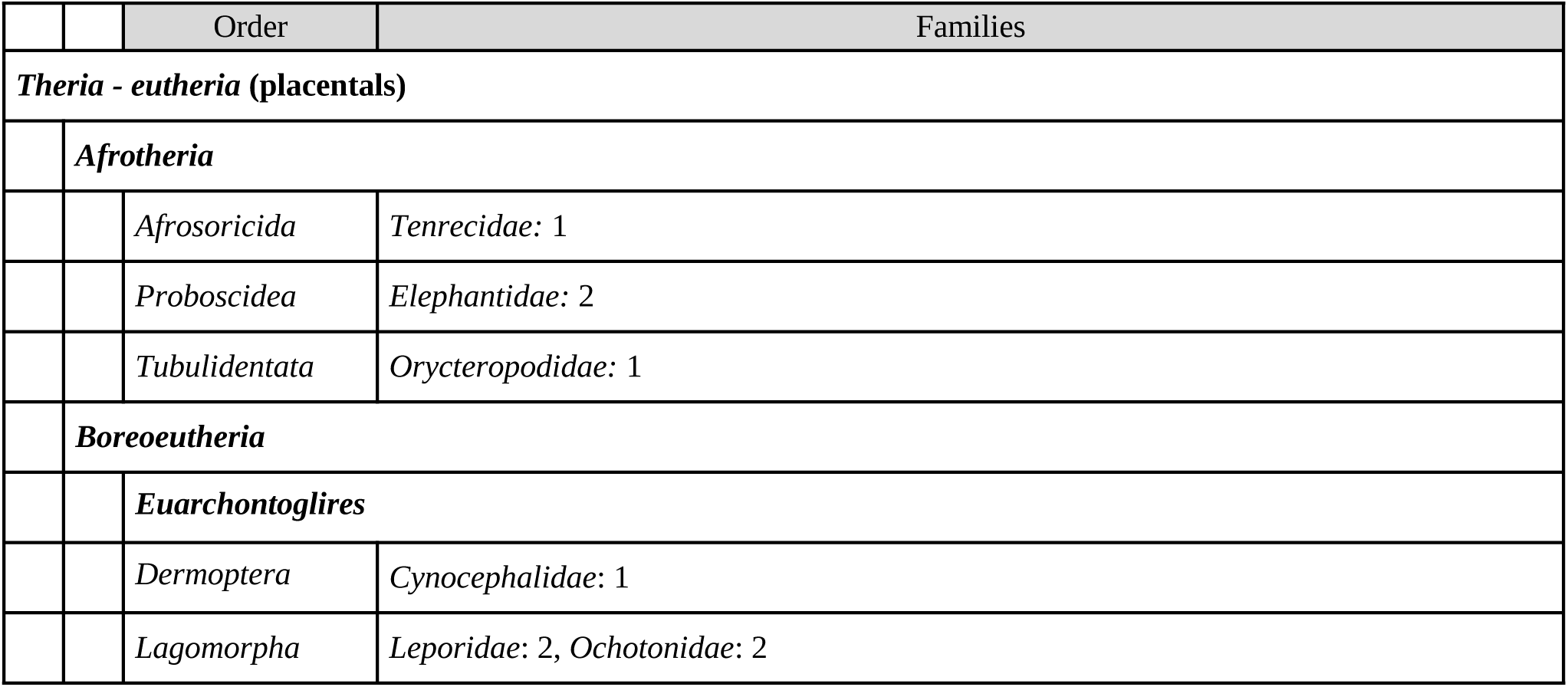
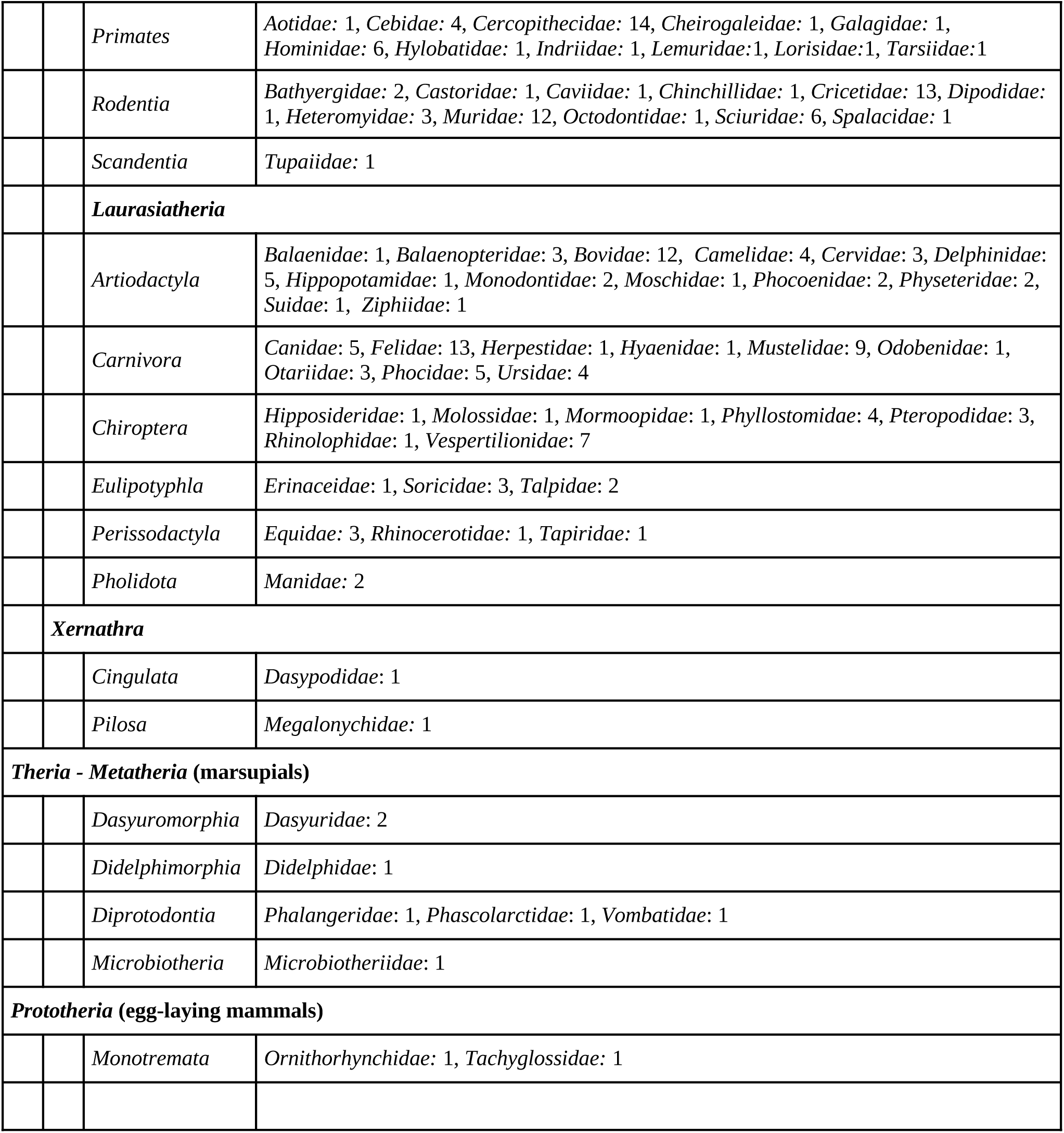
Distribution of taxa available in the mammal peptide table with COL1A1 and COL1A2 markers. Taxonomic order and family are given.

### 2.4 Taxonomic assignment with peptide tables

Peptide tables can be used to screen mass spectra and find the optimal assignment for each spectrum. The rationale is shown in Figure 2. PAMPA compares the peaks of the spectra with the masses of the peptide table and selects the more relevant taxa. This is achieved by computing a P-value score, which evaluates the significance of the match between a taxon’s peptides and the assigned spectrum. This score represents the probability of observing at least that number of matching peaks, given the total number of peaks in the spectrum, the distribution of peptide marker masses in the database, and the error margin. This approach accounts for species with varying numbers of peptide markers. PAMPA has two additional features. Firstly, it can incorporate a taxonomy, and results are interpreted within the context of that taxonomy. It means it can output classifications at different taxonomic levels (e.g. species, genus, family), depending on the resolutive power of the peptide markers. This gives the *assignment*, which is the smallest clade of the taxonomy containing all matching species. Along with the assignment, PAMPA also computes the uncertainty caused by species present in the taxonomy and for which peptide markers are missing. The *maximal clade* is the largest subtree of the taxonomy that contains only species found in the assignment. The difference between the assignment and the maximal clade comes from the fact that the database may lack representatives at certain taxonomic ranks, potentially leading to gaps in information that could affect interpretation Secondly, the program is able to construct near-optimal solutions, which allows the user to examine alternative predictions and to perform in-depth exploration of various assignment possibilities within the taxonomic space. The primary use of PAMPA is in *marker mode*, where a peptide table is provided. Alternatively, the program can also be supplied with a set of amino-acids sequences directly. In this configuration, PAMPA computes all tryptic markers (as detailed in Section 2.2) and generates a peptide table from the sequences. This is referred to as *all peptide* mode.

**Figure 2:**
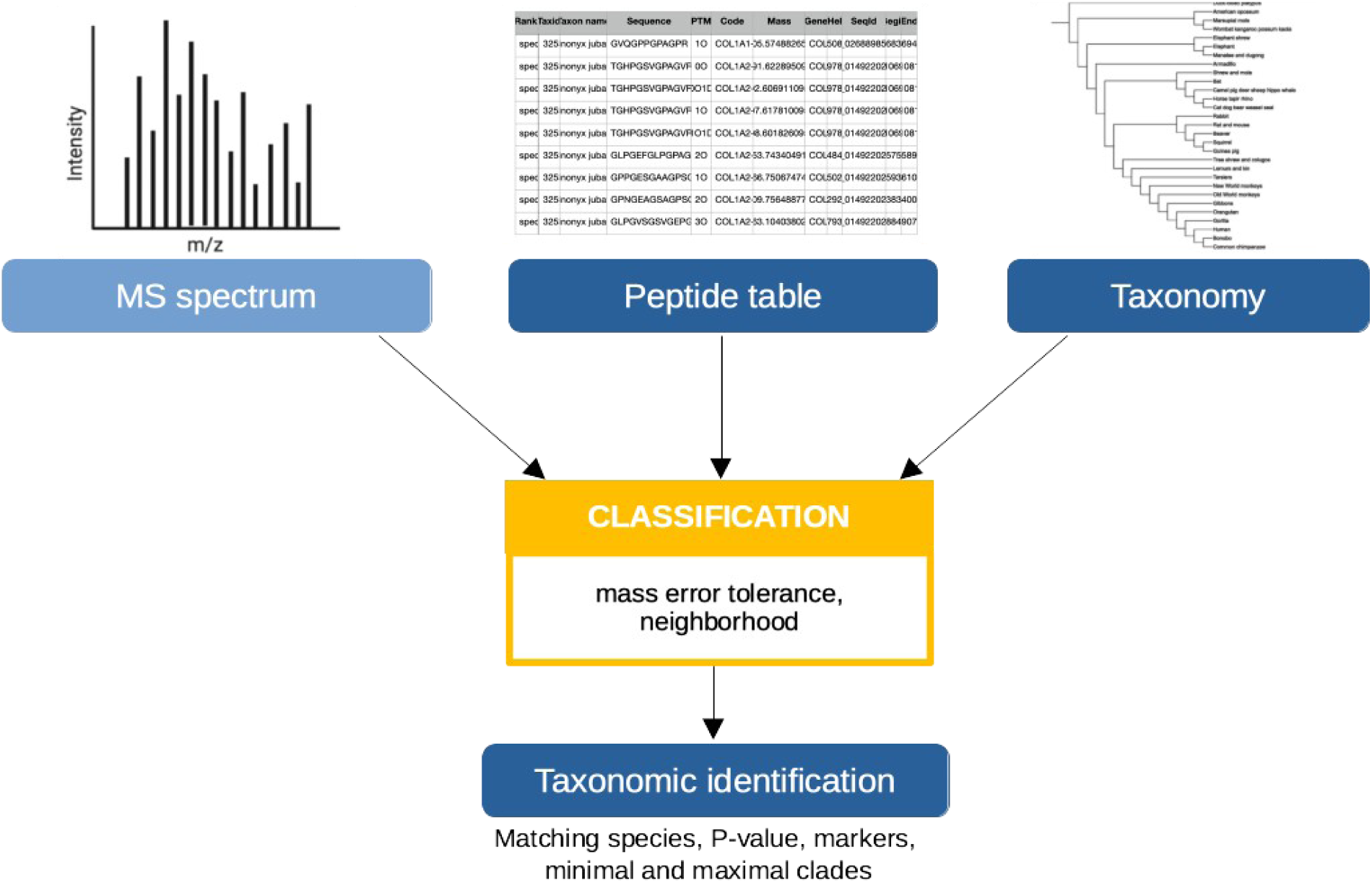
Classification of ZooMS spectra with PAMPA (marker mode). The tool takes as input a collection of MS spectra, a peptide table, and optionally, a taxonomy. It then searches for the best taxonomic classification for each spectrum among all taxa in the peptide table, based on the P-value. The output specifies which peptide markers are used to support the taxonomic prediction. The taxonomy is used to determine both the assignment and the maximal clades.

## 3. Case studies

We present the results of PAMPA using a series of examples to demonstrate the distinct functionalities of the software. Case studies 3.1, 3.2, 3.3, and 3.4 show the application of PAMPA for species identification on a variety of collections of MALDI-TOF and MALDI-FTICR spectra from various periods of time (moderns, Holocene and Pleistocene). These case studies also allowed us to compare our results with those of MASCOT, SpecieScan and Bacollite. In case study 3.5, we show how PAMPA is able to detect the presence of several species in a single sample. Case study 3.6 illustrates how to use PAMPA to infer peptide markers by homology, and lastly case study 3.7 shows how to use PAMPA to infer peptide markers directly from genomic sequences and MS spectra.

### 3.1 Taxonomic identification of MALDI-TOF and MALDI FT-ICR modern spectra

The first dataset was generated from a collection of 19 bone specimens selected for the purpose of this paper, and representing a wide taxonomic range of mammals (see Table 2). All samples are less than 100 years old, were identified by morphology, and were analyzed using both MALDI-TOF and MALDI-FTICR instruments. This gives a total of 38 spectra. 18 out of the 19 species have genomic data available, and are present in the mammal peptide table introduced in Section 2.3. The exception is *Microtus arvalis*, whose closest species in the peptide table is *Microtus oregoni,* from the same genus.

**Table 2:** List of bones database for case study on modern spectra. The table contains the common name as well as the scientific name, the family. The type of bone, the sample accession and the laboratory are given in Supplementary materials.

| Family | Common name | Scientific name |
| --- | --- | --- |
| *Bovidae* | Bison | *Bison bison bison* |
| *Bovidae* | Goat | *Capra hircus* |
| *Bovidae* | Cattle | *Bos taurus* |
| *Bovidae* | Sheep | *Ovis aries* |
| *Canidae* | Dog | *Canis lupus familiaris* |
| *Canidae* | Fox | *Vulpes vulpes* |
| *Castoridae* | Castor | *Castor Canadensis* |
| *Cervidae* | Red deer | *Cervus elaphus* |
| *Cervidae* | Fallow deer | *Dama dama* |
| *Cetacea* | Whale | *Balaenoptera (genus)* |
| *Cricetidae* | Vole | *Microtus arvalis* |
| *Equidae* | Horse | *Equus caballus* |
| *Erinaceidae* | Hedgehog | *Erinaceus europaeus* |
| *Hominidae* | Human | *Homo sapiens* |
| *Leporidae* | Rabbit | *Oryctolagus cuniculus* |
| *Muridae* | Mouse | *Mus musculus* |
| *Muridae* | Rat | *Rattus rattus* |
| *Suidae* | Pig | *Sus scrofa domesticus* |
| *Talpidae* | Mole | *Talpa (genus)* |

PAMPA was run on each of the two datasets, MALDI-TOF and MALDI-FTICR, with two different options: in *marker* mode with the full mammal peptide table, and in *all peptide* mode from the COL1A1 and COL1A2 sequences directly for the same species. The processing time of PAMPA for all mass spectra on a Dell Optiflex 7080 computer with 32 GB Ram, an Intel® Core™ i7-10700 CPU @ 2.90 GHz and 8 cores is 3 seconds for the marker mode, and 200 seconds for the all peptides mode.

#### PAMPA’s results

Table 3 illustrates the output of PAMPA for TOF spectra in the marker mode. For each spectrum, the table lists the matching species, the number of markers, the corresponding P-value, the assignment and the associated maximal clade. The matching species represent potential candidates for the spectrum, identified by having the lowest P-value, and belonging to the same assignment because they cannot be distinguished by the markers. This example illustrates the difference between the assignment and the maximal clades. For the horse spectrum, PAMPA identifies three matching species—*Equus quagga*, *E. caballus*, and *Equus asinus*—which cannot be distinguished because they share the same 11 markers present in the spectra. The software offers every possibility. The assignment is the genus *Equus.* The maximal clade is the family *Equidae*, indicating that all Equidae in the database are members of the genus *Equus*, and no other equids were tested. Similarly, for the castor spectrum, the assignment is *Castoridae*, even if only one matching species was found, *Castor canadensis*, as no other Castoridae species (e.g., *Castor fiber*) are represented in the database. Overall, Table 3 demonstrates that the results obtained by PAMPA match the morphological identifications for all mass spectra. It found a single assignment, which included the target species among the matching species. The taxonomic resolution of the classification is influenced by both the number of markers detected and the comprehensiveness of the database. All this information about undistinguishable species, minimal and maximal clades is included in the results file of PAMPA, as well as the markers occurring in the classification and the peak.

**Table 3:**
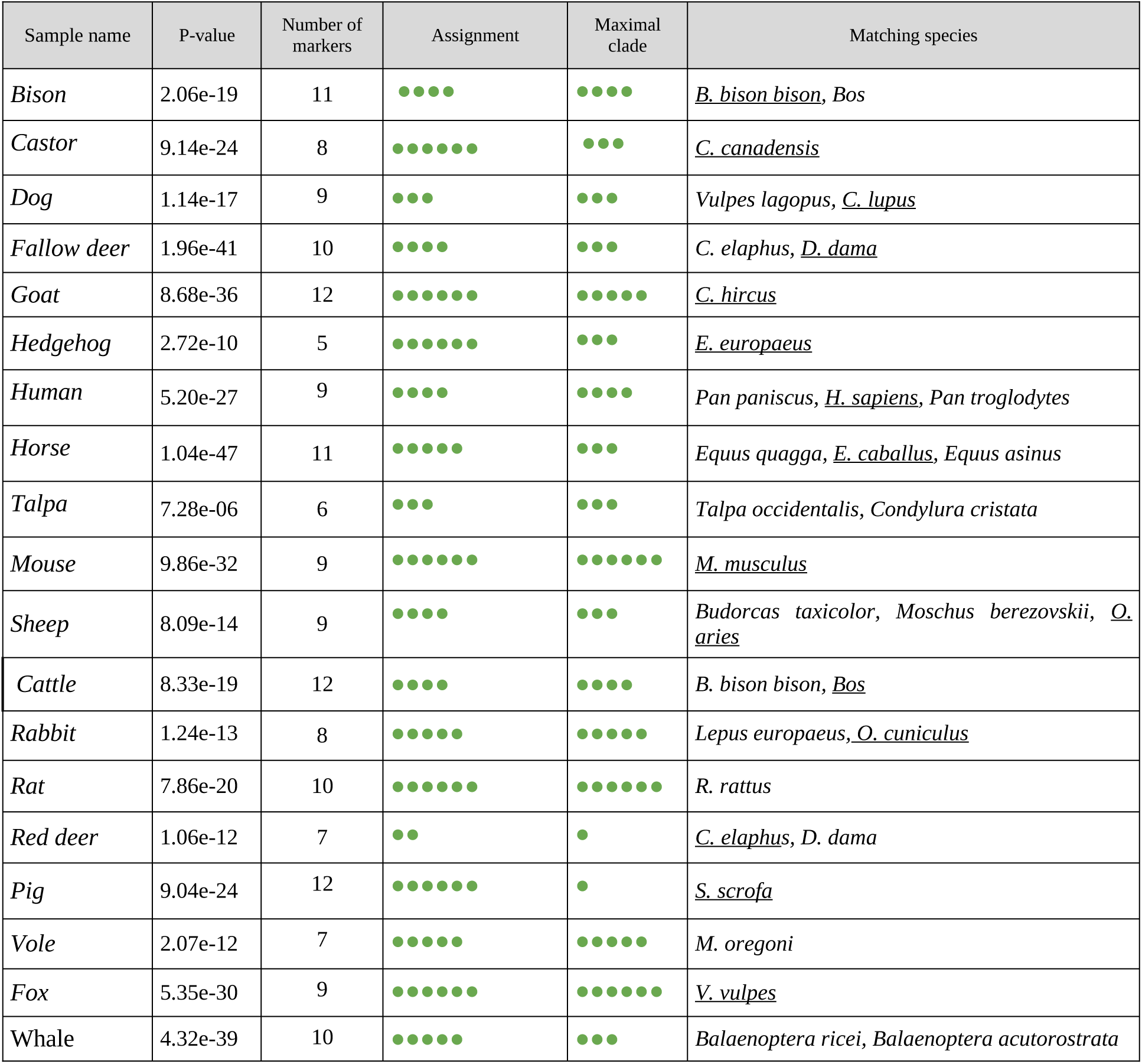
PAMPA’s results for MALDI-TOF modern spectra in the marker mode. For all spectra, the *matching species* columns show that the target species is explicitly identified as a potential match. We provide the taxonomic rank of each assignment and the maximal clade as bullet codes: ●●●●●● for species, ●●●●● for genus, ●●●● for subfamily and ●●● for family, ●● for infra-order and ● for suborder. Bos is a shortcut for Bos *indicus, Bos indicus x Bos taurus, Bos javanicus, Bos mutus, B. taurus*.

#### Comparison with MASCOT

We also ran MASCOT on this dataset. Since MASCOT is not optimized to deal with ZooMS data, we tried several settings in order to find the most suitable combination of parameters: default setting, and then with less permissive parameters (one missed cleavage, error mass tolerance 50 ppm for TOF spectra, 5 ppm for FTICR spectra), which coincides with PAMPA parameters. Since MASCOT is not intended to combine results from different proteins, we also tried it with three distinct sequence databases: COL1A1 sequences only, COL1A2 sequences only, and both COL1A1 and COL1A2 pooled together. It appeared that the best results were obtained with PAMPA parameters on the COL1A2 mature sequence database, which can be explained by the fact that COL1A2 is more variable than COL1A1. Indeed, out of the nine loci used to design peptide markers on COL1, seven come from COL1A2, and only 2 from COL1A1 (including P1 which is not variable on these species). We report here the results obtained with this setting.

Table 4 presents the complete results for all types of spectra (TOF and FTICR) in both modes (marker and all peptide), alongside the results obtained using MASCOT for comparison. In all cases, PAMPA’s results align perfectly with the known sample origins, whatever the mode. For the dog, fallow deer, and red deer spectra, the all peptide mode allowed discriminating between close species where the marker mode failed. In contrast, MASCOT produced less accurate results, particularly for TOF spectra, where only eight spectra were correctly classified. This highlights the fact that MASCOT is not well-suited for classifying MALDI-TOF spectra in ZooMS. MASCOT results improve with MALDI-FTICR spectra, because the lower mass error allows for the detection of more specific peptides. Even in this case, MASCOT still failed to consistently identify the exact target species, often detecting only closely related species.

**Table 4:**
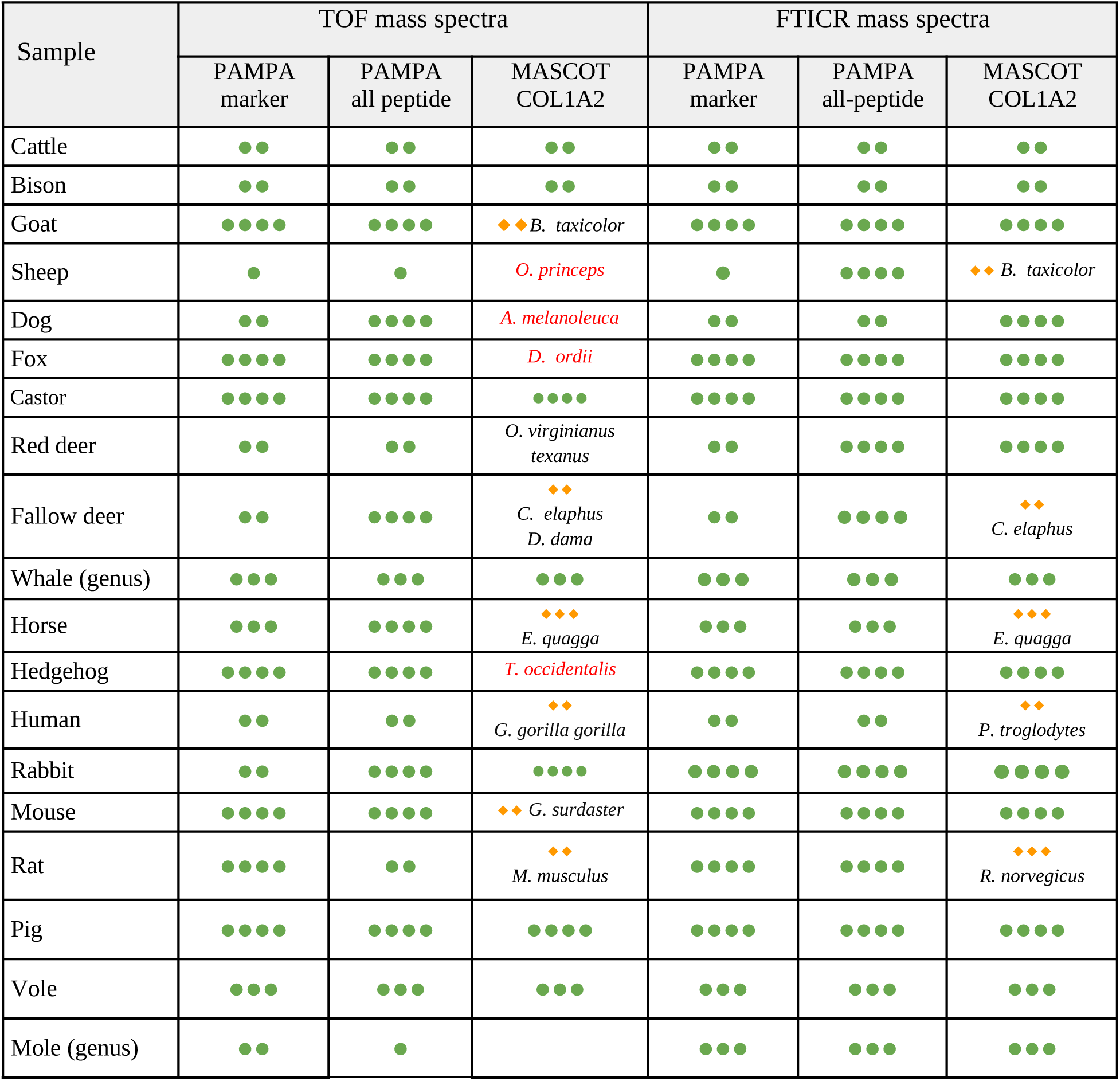
Assignments obtained with PAMPA, in marker and all peptide modes, and MASCOT for MALDI-TOF and FTICR modern spectra. The table categorizes results with ● and ◆ codes. ●●●● denotes that the identification finds a single match corresponding to the target species. ●●● (reps. ●● and ●) denotes that the target species is explicitly identified as a potential match, together with other species from the same genus (resp. family and order). ◆◆◆ (resp. ◆◆, ◆) signifies that the classification does not identify the target species, but instead finds one or more species belonging to the same genus (resp. subfamily and family). In this last case, the name of the suggested species is reported. Lastly, when no species from the same family is found, the result is reported in red.

### 3.2 Analysis of 104 archaeological MALDI FTICR mass spectra

This dataset is composed of 104 MALDI FTICR spectra coming from 104 bone samples of herbivores from the Middle Paleolithic site of Caours (Somme, France). Among these samples, 45 were identified by their osteomorphology, and taxonomic identification was conducted manually for the 59 remaining samples from the ZooMS spectra.^26^ We ran PAMPA on the whole set of spectra, and compared the identification results. Figure 3 shows the distributions of assignments obtained and the Supplementary data contains the output results. For the samples with morphology-based identification, PAMPA successfully identified all Castor, Bovinae, Sus, and Rhino specimens at the genus or subfamily level, depending on the specificity of the markers. As expected, the two non-mammalian samples (bird and turtle) were left unassigned by PAMPA, reflecting the fact that these clades are not included in the reference database proposed by the peptide table. Most *Dama dama* and *Cervus elaphus* samples were accurately identified as *Cervinae*. Two were classified as the larger infraorder Pecora (even-toed hoofed mammals with ruminant digestion), suggesting that the corresponding spectra might lack several key markers needed for more precise classification. PAMPA also failed to identify the expected rodent sample, likely due to collagen degradation or the limited representation of rodents in the database. Overall, PAMPA proved highly accurate for the large majority of samples, and in all cases does not produce any classification that is not consistent with morphological identification. As for the samples for which morphological identification was not possible, PAMPA results were in perfect alignment with the manual identifications provided in Bray *et al*., (2023), matching all assignments. Additionally, of the five manually unassigned samples, PAMPA was able to classify one as *Cervinae*, two as *Pecora*, and left only two completely unassigned. This demonstrates PAMPA’s capability to reproduce, and even improve, manual classification.

**Figure 3:**
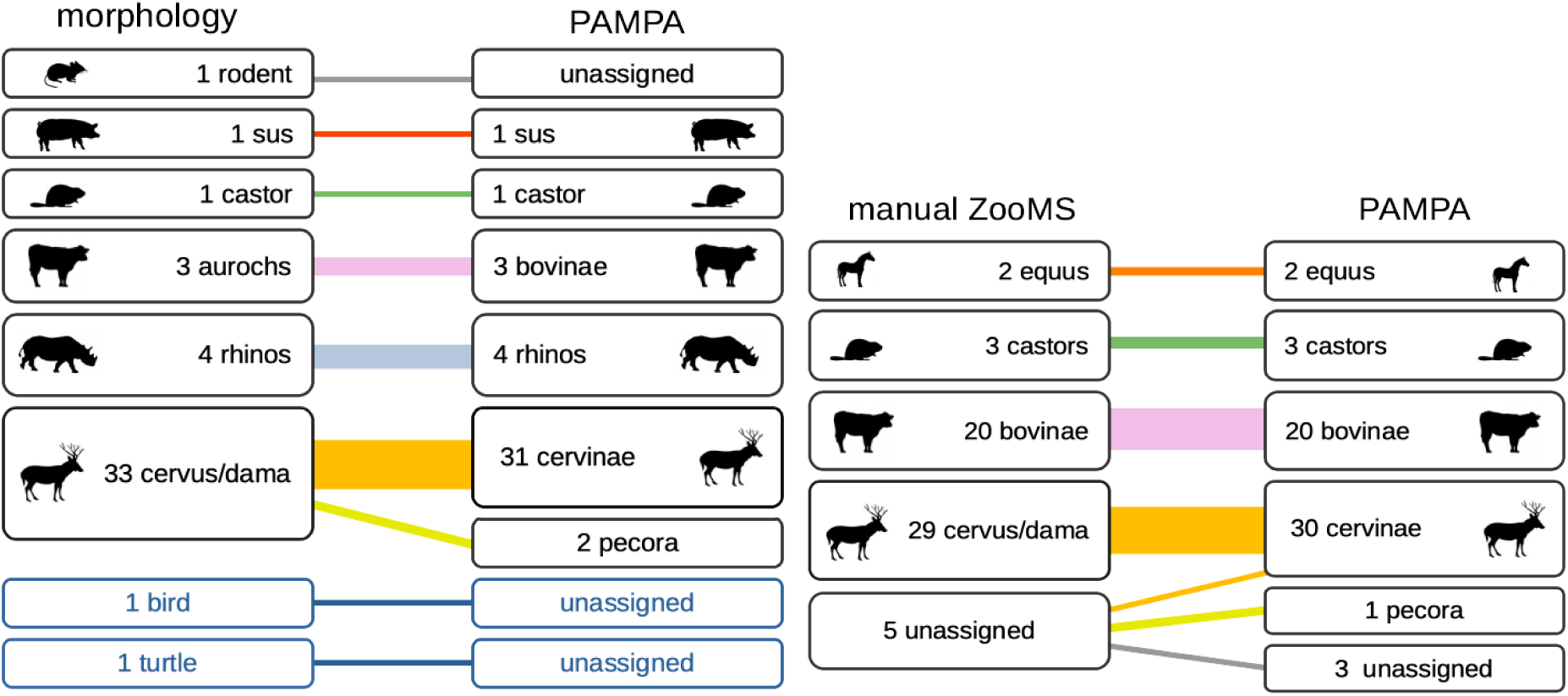
Results for the Caours case study. We show the distribution of assignments obtained with PAMPA for the 45 spectra with morphological identification (left) and for the 59 spectra with manual ZooMS identification (right).

### 3.3 Taxonomic identification of sheep and goat datasets

This case study is taken from Viñas-Caron *et al.,* and comprises 99 MALDI-TOF spectra in triplicate, obtained from 33 samples of Middle-Age parchments made of sheep ( *Ovis aries*) or goat (*Capra hircus*) skin.^31^ These data are interesting because only one peptide marker can discriminate between the two animals. This is peptide COL1A2 757-789 corresponding to the sequences GPSGEPGTAGPPGTPGPQGLLGAPGFLGLPGSR with *m/z* 3017.4963 (G) and *m/z* 3033.4912 (G’) for sheep, and GPSGEPGTAGPPGTPGPQGFLGPPGFLGLPGSR with *m/z* 3077.4963 (G) and m/z 3093.4912 (G’) for goat. We also use this example to explore the impact of preprocessing on sample classification. To do so, we apply two types of preprocessing: consensus processing, as described in Viñas-Caron *et al.,* which resulted in 33 consensus spectra, and individual analysis, where each spectrum is processed in isolation without merging or aligning it with others (see Materials and Methods). We also compared the marker mode against the all peptide mode of PAMPA. The results are shown in Figure 4. All approaches correctly classify at least 28 out of the 33 samples (84%) and 26 samples are correctly classified by all four approaches (78%). In this example, the most robust approach is the marker mode combined with individual analysis, which correctly identifies 31 out of 33 samples and fails to provide a result for two samples (ambiguous samples), with no misclassified sample. All other approaches yield two to four ambiguous samples and one misclassified sample. These results suggest that while the impact of preprocessing is small, it is still existing. Interestingly, applying a majority vote across the four approaches eliminates all misclassifications, leaving only two unclassified samples

**Figure 4:**
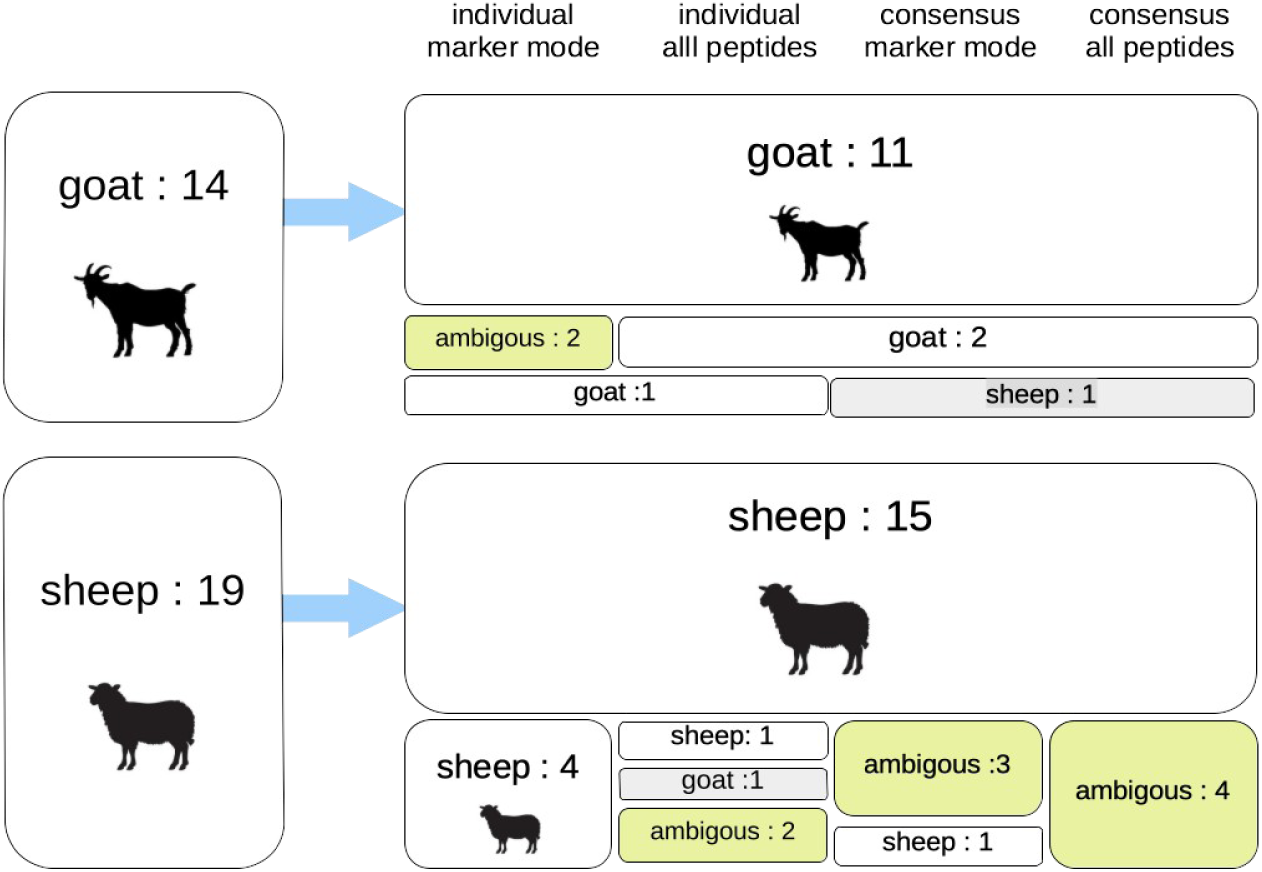
Results for the goat and sheep case study. We report results for consensus and individual spectra, in marker and all peptide modes. For individual spectra, we apply a majority rule on the three assignments provided by PAMPA.

### 3.4 Taxonomic classification of African bovids

This example, adapted from Janzen *et al*., consists of a collection of MALDI-TOF spectra from Kalundu Mound (Zambia), primarily from bovids and a few others from other mammal specimens.^8^ Manual identification is available for these specimens, with partial morphological data also provided. This dataset was previously used in Vegh *et al*., to evaluate the performance of the SpecieScan tool.^30^ To ensure a fair comparison between PAMPA and SpecieScan, we used the same species and peptide markers as SpecieScan, focusing on spectra that SpecieScan was able to successfully pre-process during the cleaning step. This resulted in a subset of 99 spectra. Outcomes of SpeciesScan and PAMPA are given in Figure 5. PAMPA achieves a higher overall percentage of correct results (90%) compared to SpecieScan (80%). Additionally, it improves over SpecieScan across all subcategories. Lastly, PAMPA provides better classifications than manual identification for 9 spectra.

**Figure 5:**
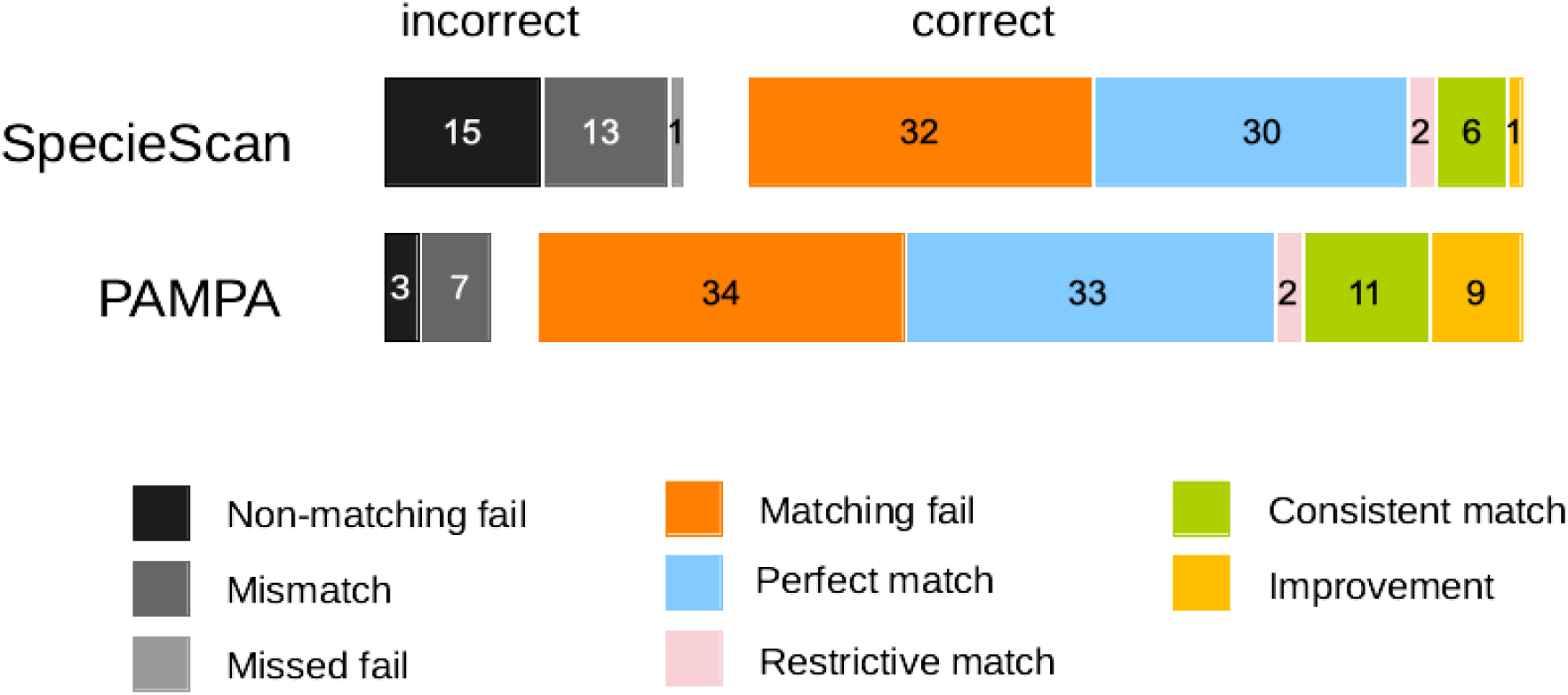
comparison of results of SpecieScan and PAMPA on the 99 MALDI-TOF African bovids spectra of Kalundu Mound. The categories (from left to right in the diagram) refer to the correct and incorrect For these 99 spectra, we classified the predictions from both SpecieScan and PAMPA as either *correct* or *incorrect*, further dividing these two categories into eight detailed subcategories. For correct results, there are five subcategories: matching fails - the tool correctly determines that the spectrum cannot be used, matching the manual analysis conclusion; perfect match - the tool’s result exactly matches the manual analysis; restrictive match - the tool’s result is a subset of the manual analysis outcome, meaning the tool’s result is included within the manually obtained result; consistent match - the tool’s result encompasses the manual analysis outcome, meaning it includes the manual analysis; improvement - the tool’s result, while included in the manual analysis outcome, is particularly consistent with morphological observations, suggesting an enhancement over the manual classification. For incorrect results, we identify three subcategories: non-matching fails -the tool fails to produce any result, even though manual analysis does provide an answer; mismatches - the tool’s prediction does not align with the result from manual analysis; missed fails - the tool provides a result, but this is an error, as the MS spectrum actually contains no useful information.

To go deeper into the analysis, we focused on spectra with full identification—those classified either manually or morphologically at the genus level for bovids, and the family level for other mammals. This narrowed our analysis to a set of 50 spectra. The comparative results of PAMPA and SpecieScan are shown in Figure 6. PAMPA produced 44 correct results, including 29 perfect matches, 10 consistent matches, and five improved classifications over manual classification. There were only six incorrect results: three spectra with no outcome and three spectra incorrectly assigned to *Bos* (for which only morphological identification was possible). In contrast, SpecieScan yielded 31 correct results (26 perfect matches and five consistent matches) and 19 incorrect results (10 fails and nine mismatches).

**Figure 6:**
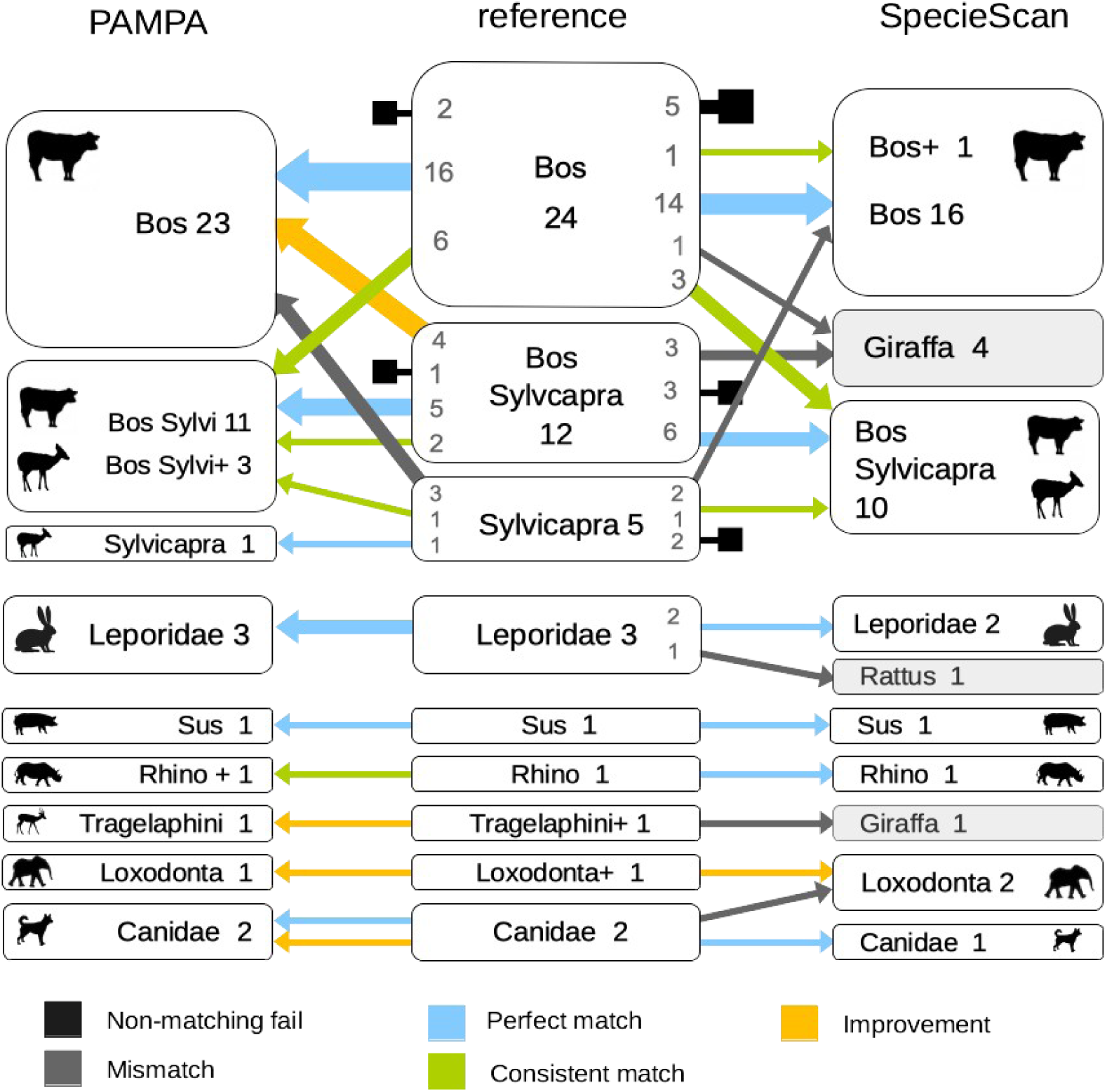
Results for African bovid spectra with full identification. The color code is the same as for Figure 5. Groups with the symbol “+” contain the designated species, as well as other species. Taxa found by SpecieScan that are not expected to be found in these samples (*Giraffa* and *Rattus*) are highlighted in gray.

### 3.5 Detection of mixtures

Contamination between multiple bones may occur during sampling, posing a challenge for species identification. To simulate this scenario, we created mixed datasets by performing MALDI-FTICR digestions on bone powder mixtures from two species, each conducted in duplicate. The mixtures analyzed are *Capra hircus* with *Bos taurus*, *Ovis aries* with *Capra hircus* and *Ovis aries* with *Cervus elaphus*. These pairs of species were chosen because they share common peptide markers. PAMPA software was tested on these spectra using the “neighboring” option, that generates all near-optimal solutions. We chose a threshold of 85%, meaning that all assignments with a number of recognized markers at least equal to 85% of the number of markers in the optimal solution are retained. PAMPA correctly identifies two clades for each of the three samples (Supplementary Material).

### 3.6 Generation of new peptide markers by homology

In this case study, we illustrate the usage of PAMPA’s homology module on a selection of marine mammals. As of now, 25 species have at least one COL1A1 sequence and one COL1A2 sequence available in the NCBI Protein database for those taxonomic groups, among them 20 have proteomic data Buckley *et al.,* (2014), Hofman *et al.*, (2018). ^32, 33^ All together, these species cover a large taxonomic diversity, composed of 15 cetaceans, with 4 mysticeti (baleen whales) and 12 odonceti (tooth whales), and 9 pinnipeds. The list is given in Table 5.

**Table 5:**
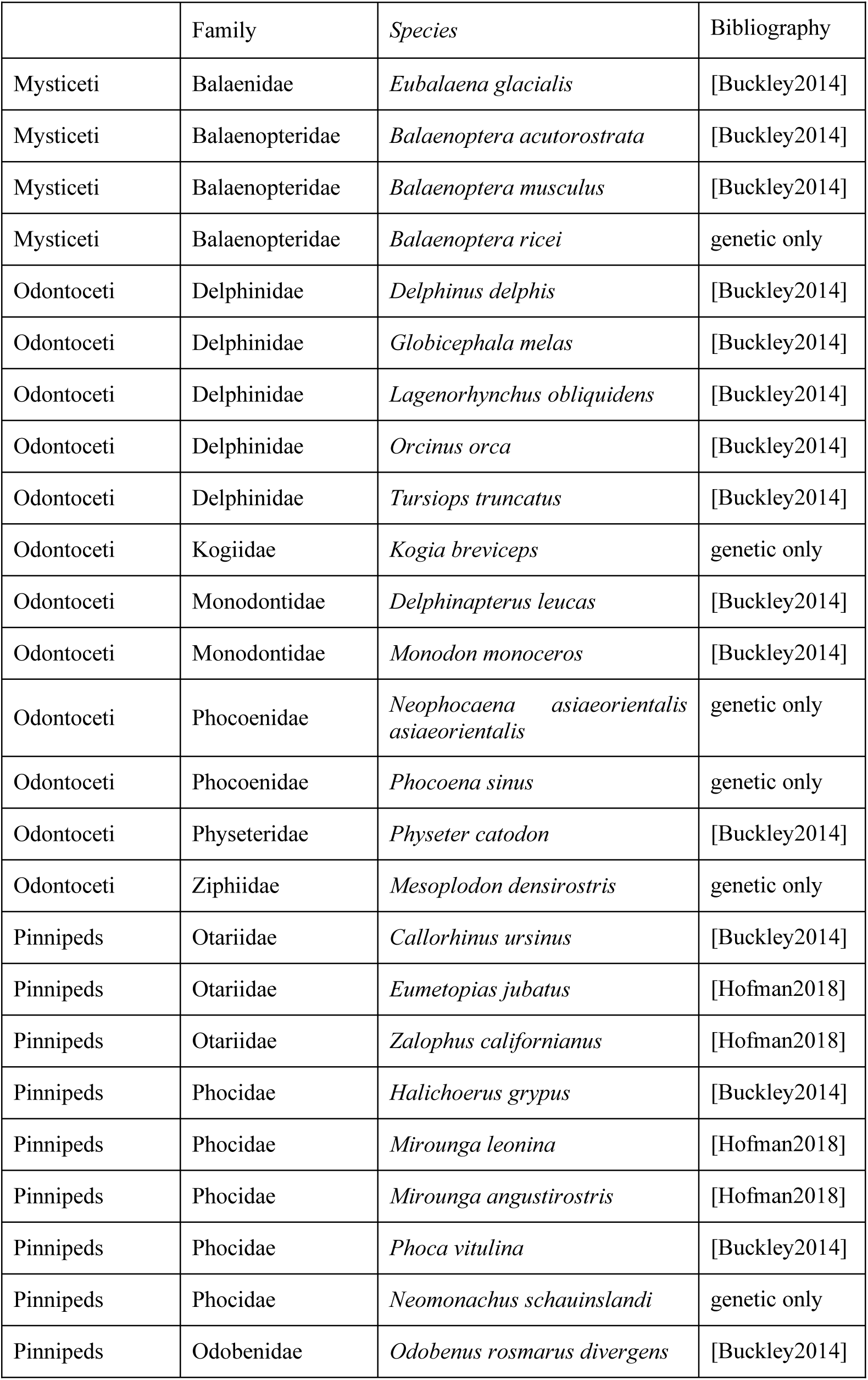
List of marine species used in the homology search case study.

We selected three species to serve as references for the homology search: *T. truncata* (odonceti) and *B. musculus* (mysticeti) for cetaceans with peptides Cet1, Cet2, A, B, C, D, F and G on the one hand, and *O. rosmarus* for pinnipeds with peptides Cet1, P, A, B, C, D, F, G on the other hand. Using these references, PAMPA automatically identified the markers present in the COL1A1 and COL1A2 sequences of the 22 remaining species. Results are visible on Table 6 for pinnipeds and Table 7 for cetaceans. Buckley *et al.,* (2014) also provided LC-MS/MS sequences for marker D for 15 species: *O. rosmarus, C. ursinus, P. vitulina, H. grypus, T. truncatus, D. delphis, L. obliquidens, G. melas, O. orca, M. monoceros, D. leucas, P. catodon, E. glacialis, B. musculus, B. acutorostrata.* We present in Table 8 the sequences found by PAMPA for this marker. The full peptides tables, with sequences, position and PTMs are available in Supplementary material.

**Table 6:**
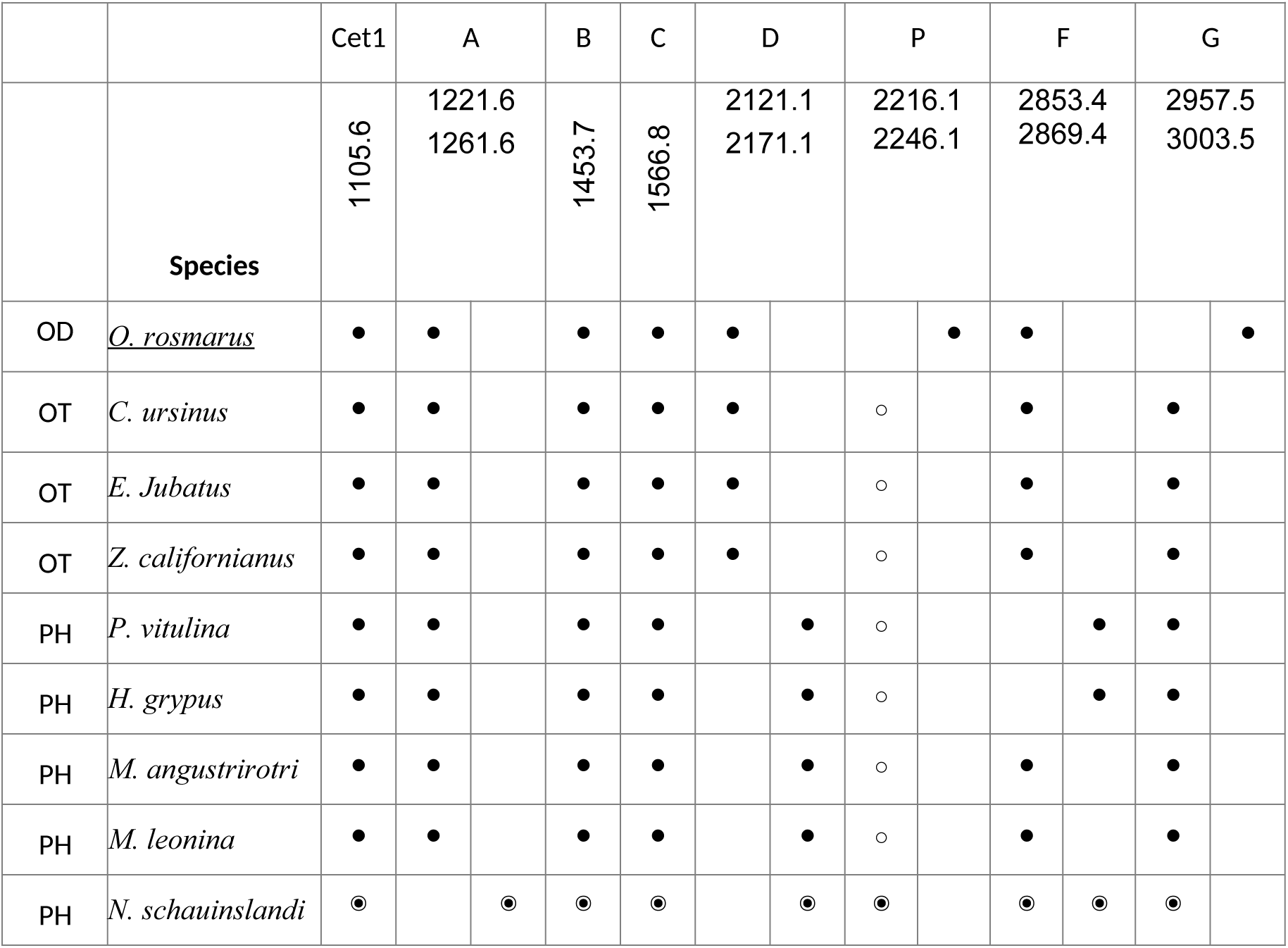
Homology results for markers Cet1, P, A, B, C, D, F, G for pinnipeds. Peptides of *O. rosmarus* (underlined) are used as reference. Results for species without available proteomic data are marked with ◉. For other species, *m/z* values that match the literature are represented by ⚫, while values that do not align with the literature are indicated with ⚪. The first column is for the family: OD for *Odobenidae*, OT for *Otariidae* and PH for *Phocidae*.

**Table 7:**
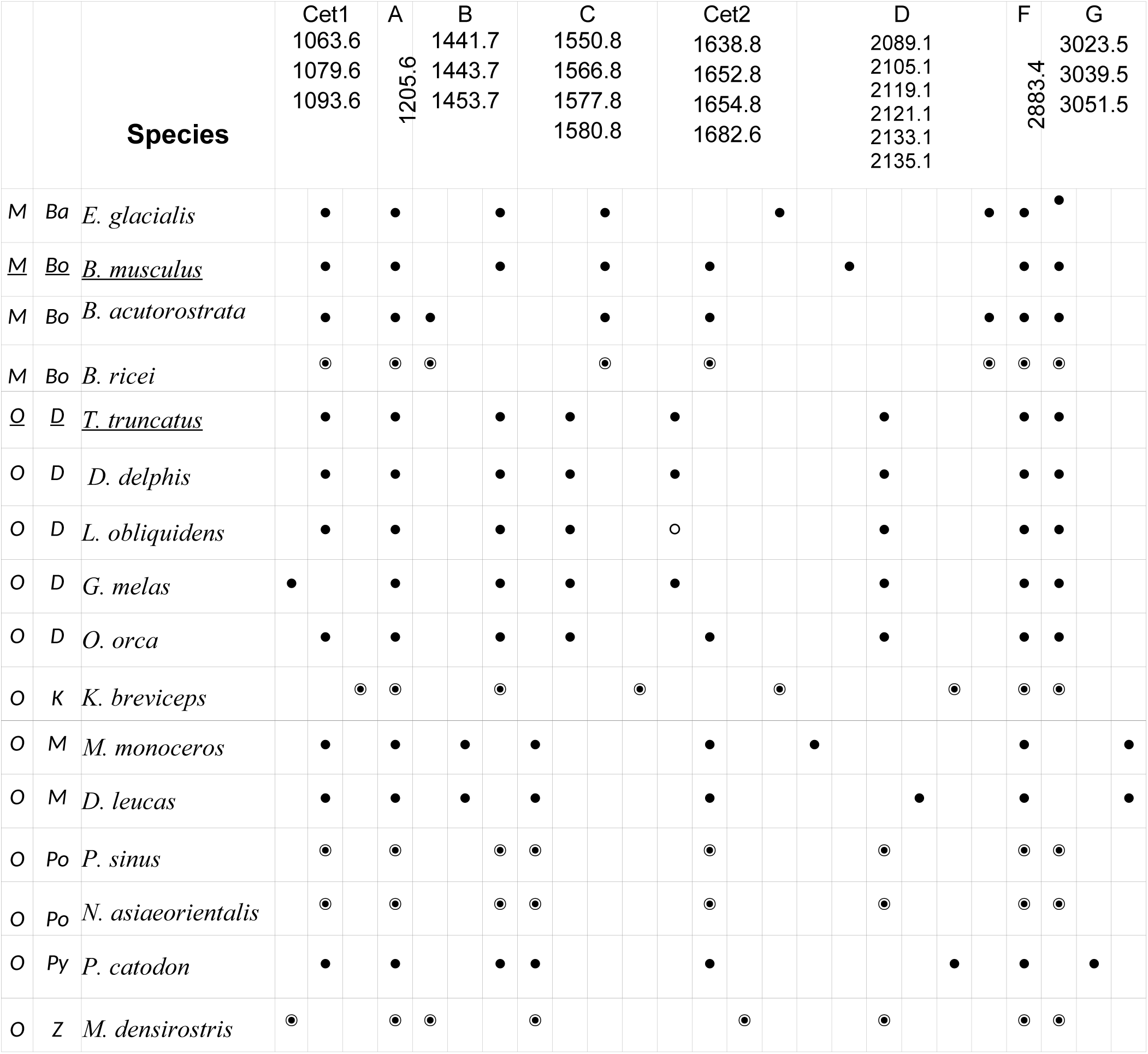
Homology results for markers Cet1, Cet2, A, B, C, D, F, G for *cetacea* (odontoceti and mysticeti). Peptides of *T.truncata* and *B. musculus* (underlined) are used as reference. Results for species without available proteomic data are marked with ◉. For other species, *m/z* values that match the literature are represented by ⚫, while values that do not align with the literature are indicated with ⚪. The first column displays the parvorder: M for mysticeti, and O for odotonceti, the second the family: Ba for *Balaenidae*, Bo for *Balaenopteridae*, D for *Delphinidae*, K for *Kogiidae*, M for *Monodontidae*, Po for *Phocoenidae*, Py for *Physeteridae*, Z for *Ziphiidae*.

**Table 8:**
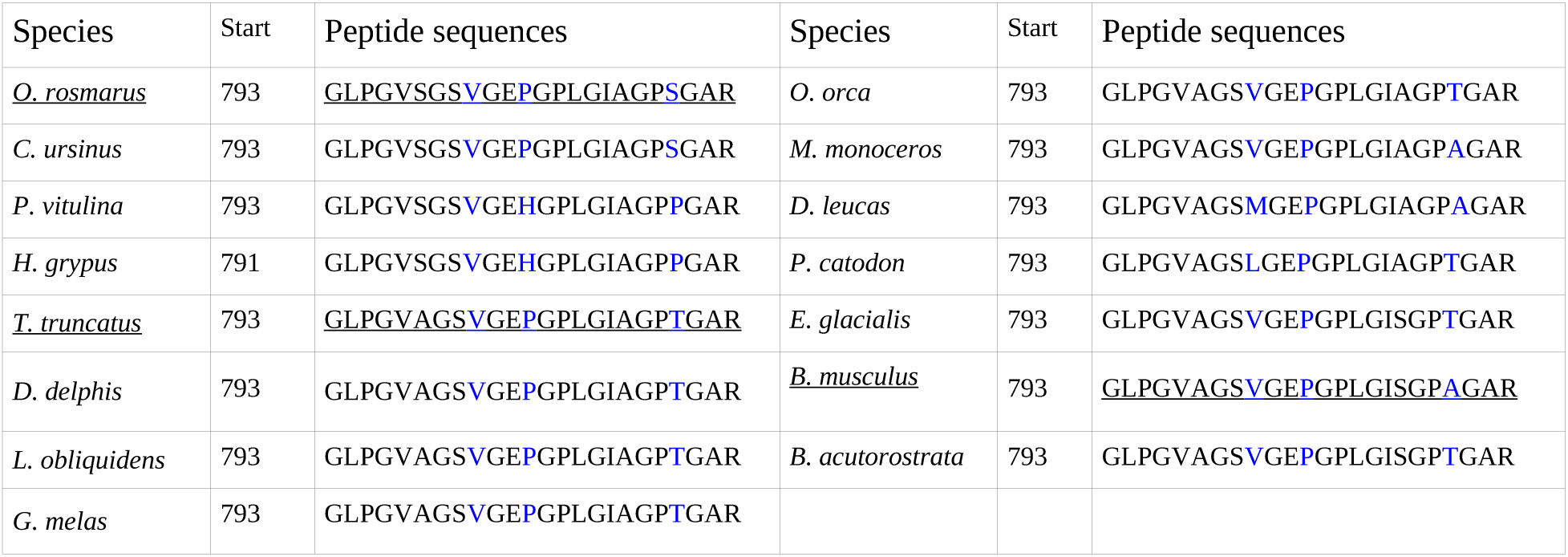
Sequences found by PAMPA for D peptide marker (COL1A2 793-816). Sequences used as models are underlined. Variable positions are in blue. The start column is for the start position within the helical region (automatically inferred by PAMPA). For all species, peptide D is at position 793, except for *H. grypus,* which here alerts on the fact that the COL1A2 *H. grypus* sequence has a poor quality.

| Species | Start | Peptide sequences | Species | Start | Peptide sequences |
| --- | --- | --- | --- | --- | --- |
| *O. rosmarus* | 793 | <u>GLPGVSGSVGE**P**GPLGIAG**P**SGAR</u> | *O. orca* | 793 | GLPGVAGSVGE**P**GPLGIAG**P**TGAR |
| *C. ursinus* | 793 | GLPGVSGSVGE**P**GPLGIAG**P**SGAR | *M. monoceros* | 793 | GLPGVAGSVGE**P**GPLGIAG**P**AGAR |
| *P. vitulina* | 793 | GLPGVSGSVGE**H**GPLGIAG**P**PGAR | *D. leucas* | 793 | GLPGVAGSMGE**P**GPLGIAG**P**AGAR |
| *H. grypus* | 791 | GLPGVSGSVGE**H**GPLGIAG**P**PGAR | *P. catodon* | 793 | GLPGVAGSLGE**P**GPLGIAG**P**TGAR |
| *T. truncatus* | 793 | <u>GLPGVAGSVGE**P**GPLGIAG**P**TGAR</u> | *E. glacialis* | 793 | GLPGVAGSVGE**P**GPLGISG**P**TGAR |
| *D. delphis* | 793 | GLPGVAGSVGE**P**GPLGIAG**P**TGAR | *B. musculus* | 793 | <u>GLPGVAGSVGE**P**GPLGISG**P**AGAR</u> |
| *L. obliquidens* | 793 | GLPGVAGSVGE**P**GPLGIAG**P**TGAR | *B. acutorostrata* | 793 | GLPGVAGSVGE**P**GPLGISG**P**TGAR |
| *G. melas* | 793 | GLPGVAGSVGE**P**GPLGIAG**P**TGAR |  |  |  |

For pinnipeds, the theoretical masses associated to the found peptides are all consistent with those reported in Buckley *et al*., and, Hofman *et al*., except for marker P in Otariidae and Phocidae (marked with ⚪ in the table).^32, 33^ For this marker, PAMPA identifies the peptide sequence GETGPAGRPGEVGPPGPPGPAGEK at the correct helical position. The calculated mass (with three hydroxyprolines) is 2216.05, whereas the reported mass these publications is *m/z* 2215. However, the MALDI spectra provided as evidence for *m/z* 2215 raise uncertainty. Several spectral data set actually show a peak at 2216, annotated as P: Figure 4 for the genus Phoca, supplementary Figure 9 in Buckley *et al*., (2014) for *Callorhinus ursinus*, Otariidae spectra and Mirounga spectra in Figures S5 and S8 in Hofman *et al*. (2018) Note that the confusion is unlikely to result from an additional deamidation, as the peptide sequence is not susceptible to deamidation.

For cetaceas, values found by PAMPA also globally match the expected values. The only exception is the mass of the Cet2 marker for *L. obliquidens*, which is reported to be 1652 and for which PAMPA predicts *m/z* 1638.8. However, a closer look at the supplementary material of Buckley *et al*., (2014) shows that the MALDI-TOF spectrum for *L. obliquidens* is missing, which raises concerns regarding the accuracy of the value for Cet2. Moreover, the *m/z* Cet2 value for *Lagenorhynchus albirostris*, belonging to the same genus is shown to be *m/z* 1638.8 in the same article. Here again, these two pieces of information taken together make PAMPA’s prediction plausible.^33^

We also aimed to assess the impact of the new species on the diagnostic power of peptides. Markers Cet1 and A are known to differentiate cetaceans from pinnipeds, and the newly observed *m/z* values support this distinction. Peptide A is at *m/z* 1205 in cetaceans, while in pinnipeds, it appears at *m/z* 1221 and 1261. Peptide Cet1 shares the value *m/z* 1105 with the large majority of mammals, while cetaceans exhibit a series of alternative *m/z*, with a new modality (1093) for Kogiidae. Within pinnipeds, the addition of *N. schauinslandi* (Phocidae) does not alter the classification of the three pinniped families. Odobenidae, Otariidae, and Phocidae can still be distinguished based on a combination of peptide markers D and G: peptide D has an *m/z* 2121 in Odobenidae and Otariidae but shifts to *m/z* 2171 in Phocidae, while Peptide G is *m/z* 3003 in Odobenidae and *m/z* 2957 in Otariidae and Phocidae. Among cetaceans, the addition of five new species adds complexity to the picture. The two newly analyzed Phocidae species share markers with both Monodontidae (markers Cet1, Cet2, C and G) and Delphinidae (markers B and D). Meanwhile, *K. breviceps* and *M. densirostris* introduce new variants for peptides Cet1, Cet2 and C, further highlighting the genetic diversity of cetaceans. Of course, all these observations come only from genetic information and should be confirmed by proteomics.

Regarding the peptide sequences for marker D (Table 8), they all match the LC-MS/MS sequences with 100% identity. Here PAMPA is able to recover the correct homolog peptide sequence with up to 3 amino-acids substitutions.

### 3.7 De novo full peptide generation and filtering

In this last case study, we illustrate the application of de novo full peptide generation and filtering functionality of PAMPA on COL1A1 and COL1A2 sequences from *O. aries* (XP_027830506.1 and XP_004007775.1). On these two sequences, the all peptide option of PAMPA generated a total of 185 tryptic peptides, allowing for up to one missed cleavage. Incorporating the presence of hydroxyprolines into these peptides yielded 228 distinct *m/z* values. We then performed peak filtering by running PAMPA on the same sequences together with all the individual spectra of case study 3.4 coming from the 19 samples identified as *O. aries* by the authors of [Viñas-Caron2023]. We obtained a refined set of peptides, consisting of 49 peptides with 71 masses (including hydroxylation and deamidation). This subset includes 9 out of the 12 ZooMS peptide markers, which all have correct *m/z* and helical position: COL1A1 508-519 (P1), COL1A2 978-990 (A and A’), COL1A2 484-498 (B), COL1A2 502-519 (C), COL1A2 292-309 (P2), COL1A2 793 - 816 (D), COL1A1 586-618 (F), COL1A2 757-789 (G’). Only peptides E, F’ and G, are missing, which is consistent with the observation that they are the first to be lost as part of collagen degradation and are often the hardest to identify.^8, 34, 35^ Additionally, peptide COL1A2 889-906, which has been previously reported in other bovidae, is observed in 95% of spectra.

## 4. Materials and Methods

### Development of PAMPA

The software is written in Python 3.7 and necessitates two libraries: Biopython and pyteomics. It consists of two primary scripts: PAMPA CLASSIFY that performs species identification from peptide tables and taxonomies, and PAMPA CRAFT that enables the user to define their own markers by creating custom peptide tables.

### Preparation of MALDI-TOF and MALDI FT-ICR modern spectra dataset (case study 3.1) and “mixtures” (case study 3.5)

#### Bone specimens

Bone specimens come from collection from the labs EEP (UMR 8198, Evolution, Ecology and Paleontology), HALMA (UMR 8164, Histoire, Archéologie et Littérature des Mondes Anciens) and MSAP (UAR 3290 - MSAP - Miniaturisation pour la Synthèse, l’Analyse et la Protéomique) at University of Lille.

#### Mixture preparation

A mixture of 1 mg of bone powder from each species was placed in an eppendorf^Ⓡ^ 1.5 mL. The mixtures analyzed are *Capra hircus* with *Bos taurus*, *Ovis aries* with *Capra hircus* and *Ovis aries* with *Cervus elaphus*.

#### Digestion of bones

Proteomics samples were generated specifically for this study. The digestion of bones follows the same protocol as the one described in Bray *et al* (2023).^26^ Briefly, 1 mg of bones was deposited in 96 wells plate MultiScreenHTS-IP, 0.45 µm (MSIPS4510, Millipore, Billerica, MA, USA). The bone powder was demineralized, washed, gelatinized and digested in the 96 wells plate MultiScreenHTS-IP. The demineralization solution was digested in classical 96 well plates. The peptides form bones and demineralized solution were purified on C18 96 well plates (Affinisep, Petit-Couronne, France).

#### Purification of peptides

Briefly, the plate was washed once with 500 µL of acetonitrile (ACN) followed by a washing step with 80% ACN, H_2_O 0.5% acetic acid repeated 3 times and a second washing repeated 3 times with H_2_O alone 0.5% acetic acid. Tryptic peptides from bone powder were resuspended in 200 µL of a H_2_O, 0.5% acetic acid solution. Tryptic peptides form both solutions were transferred to C18 96-well plate and eluted with a vacuum manifold. The plate was washed 3 times with 200 µL of H_2_O, 0.5% acetic acid. Peptides were recovered in a V-bottom well collecting plate using 100 µL of 80% ACN, 0.1% acetic acid solution followed by 100 µL of ACN. The plate was evaporated on TurboVap 96 Evaporator (Caliper LifeScience, Hopkinton, USA). For mass spectrometry analysis, the sample was dissolved again in 10 μl of H_2_O 0.1% formic acid.

#### Mass spectrometry analysis

Desalted peptides (1 µL) were deposited on 384 Ground steel MALDI plates (Bruker Daltonics, Bremen, Germany) or 384-well AB Sciex MALDI plates, then 1 µL of HCCA matrix at 10 mg/mL in ACN/H_2_O 80:20 v/v 0.1% TFA was added for each sample spot and dried at ambient temperature. MALDI TOF mass spectrometry was performed on an AB Sciex 4800 Plus MALDI TOF/TOF (AB Sciex, Foster City, CA, USA) equipped with a 355 nm solid-state laser and using AB Sciex 4000 Series Data Explorer control and processing software. Two thousand shots per spectra were accumulated in the *m/z* range from 693.01 to 5,000. MALDI FTICR experiments were carried out on a Bruker 9.4 Tesla SolariX XR FTICR mass spectrometer controlled by FTMSControl software and equipped with a CombiSource and a ParaCell (Bruker Daltonics, Bremen, Germany). A Bruker Smartbeam-II Laser System was used for irradiation at a frequency of 1,000 Hz and using the “Minimum” predefined shot pattern. MALDI FTICR spectra were generated from 500 laser shots in the *m/z* range from 693.01 to 5,000 with 2 M data points (transient length of 5.0332 s). Twenty spectra were averaged. The transfer time of the ICR cell was set to 1.2 ms and the quadrupole mass filter operating in RF-only mode was set at *m/z* 600.

#### Data treatment for MALDI-TOF spectra

raw MS files were transformed into MzXml by TD2converter, processed with mMass v5.5.0.^36^ for deisotoping, and calibrated with ZooMS peptide list. Baseline correction was applied with precision to 15 and relative offset to 25. Peaks were detected with a S/N (signal-to-noise ratio) > 3, quality 0.5. Deisotoping step was realized with charge1, isotope mass tolerance 0.1, isotope intensity tolerance 25%, label envelope tools 1st envelope, envelope intensity envelope maximum. Lastly, csv files were created for treatment with MASCOT and PAMPA.

#### Data treatment for MALDI-FTICR spectra

raw MS files were processed using Compass DataAnalysis 5.0. Mass spectra were calibrated with the ZooMS peptide list. The SNAP™ algorithm was employed with the following parameters of S/N > 3 and quality 0.6. Automated processing was realized with a VBA script from Bruker. The list of *m/z* (mass over charge) and Intensity was exported in a csv file, and this file was directly provided to PAMPA and MASCOT.

#### PAMPA

The software was launched with recommended mass error margin values for MALDI-TOF spectra (50 ppm) and MALDI-FTICR spectra (5 ppm) respectively. Taxonomy was downloaded from NCBI and the peptide table used is the mammal peptide table described in Section 2.3.^37^

#### MASCOT search

we used MASCOT version 2.8.3 using the following parameters: enzyme = trypsin (allowing for cleavage before proline); maximum missed cleavages = 1; variable modifications = oxidation of proline; product mass tolerance for FTICR = 5 ppm and 50 ppm for MALDI TOF. The used database contains the COL1A1 and COL1A2 sequences of the species contained in the PAMPA peptide table. The MS errors used were the same as PAMPA.

### Preparation of the Caours archeological dataset (case study 3.2)

Raw MALDI-FTICR files are available from the PRIDE database with accession number PXD038283. Files were processed with the same workflow as MALDI FTICR modern spectra, and PAMPA was launched with the same parameters. The only difference is that we allowed for deamidation, given that we are working with ancient samples.

### Preparation of the Sheep and Goats dataset (case study 3.3)

Raw data for MS spectra were retrieved from https://zenodo.org/records/7469803. We only kept MS spectra for which triplicates were available (99 spectra). For the consensus spectra, MzML files were processed with the R script processing.md used in [Hickinbotham202]. Doing that, we obtained 33 consensus spectra. For the individual analysis, files were processed with the same workflow as MALDI TOF modern spectra, described in case study 3.1. PAMPA was launched with peptide markers for *Ovis aries* and *Capra hircus*. Deamidation was allowed.

### Preparation of the Zambia dataset (case study 3.4)

MS raw spectra were downloaded from https://zenodo.org/records/3971142. We selected all spectra with prefix KAV, which correspond to the Kalundu Mound site. These spectra were processed using the preprocessing code of SpecieScan.^30^ For that, the signal-to-noise ratio was set to 3 and the parameters of the monoisotopicPeaks function were set minCor: 0.90, tolerance: 1e-4, distance: 1.00235 and size: 2L:10L. The technical replicate spectra were combined, and a total of 99 spectra were obtained. For the database of peptide markers, we used the same list of species and markers as SpecieScan (supplementary material) and retrieved the corresponding masses from Presslee compilation.

## 4. Discussion

Case studies 3.1-3.5 show that PAMPA was able to successfully perform taxonomic assignment for MALDI TOF and MALDI FTICR spectra for ZooMS experiments. All results obtained through several examples and different scenarios show high reliability for identification using both data created for this study and data from published sources. In all cases, PAMPA accelerates the analysis of ZooMS spectra data and enhances reproducibility. Another distinctive feature of PAMPA is that it can be used either in basic mode or in advanced mode. The basic mode provides the user with a precomputed list of peptide markers, taxonomy and default parameter values. This is the mode we used in case studies 3.1, 3.2, 3.3, 3.5. In the advanced mode, the software is highly parameterized and flexible, allowing users to customize various settings to suit their specific needs. As an example, we generated a custom table for African Bovids in case study 3.4. The user can also choose to allow deamidations, or not, and to select which markers can undergo such PTM. Lastly, PAMPA comes with a config file that allows to specify the minimal and maximal length of peptides, the number of missed cleavages as well as the enzyme used for digestion. With these different levels of setting, we aimed to balance customization with ease of use to adapt to various needs. All examples presented in the article focus on type I collagen peptide markers, but the method has the potential to be applied to other molecules, as long as peptide markers can be identified.

Apart from species identification, PAMPA provides a range of complementary functionalities for handling marker peptides: finding them in genetic sequences, inferring them by homology, supplementing existing databases. In this perspective, case study 3.6 shows that PAMPA is able to correctly recover peptide markers by homology from the sequence alone, with no prior sequence alignment. In the example, it proves to be useful to control the quality of previously published results, question them and eventually correct them.

To enrich this discussion, we would like to highlight some limitations of PAMPA and show some directions for future research. First, PAMPA results still depend on the pre-processing of the spectra. The identification of monoisotopic peaks can be limiting, as most methods use the isotopic profile of peptides to detect the monoisotopic peak. If this profile has a low intensity, it is possible that only two peaks in the profile are present. Under these conditions, the pre-processing software will not detect the monoisotopic peak. Currently, in FTICR, monoisotopic peaks are detected using the SNAP algorithm, which is integrated into Bruker’s data analysis software. Other methods for peaks picking specialized in ZooMS are available such as Hickinbotham *et al*., or SpeciesScan. ^29, 30^ In case study 3.3, we tried to evaluate the impact of this processing step on the final result for taxonomic classification. It would be interesting to expand this analysis by incorporating multiple sample sources and diverse preprocessing methods. Second, some alternative bioinformatic approaches employ machine learning methods and supervised classification to analyze mass spectra without any prior information on peptide markers.^38, 39^ They compare peaks with a pre-existing database of annotated MS spectra. Although potent, such methods necessitate a significant number of precisely labeled mass spectra generated under similar conditions to mitigate the impact of artifacts, variations, and discrepancies, such as contamination. It could be a fruitful avenue of research to explore how to automatically combine partial knowledge on peptide markers and such approaches.

## 5. Conclusion

We have introduced a new bioinformatics tool, called PAMPA, specifically designed to assist in processing MALDI TOF and MALDI FTICR spectra for ZooMS experiments. PAMPA provides a range of complementary functionalities for handling peptide markers and performing taxonomic assignments. It has been designed to be used by researchers with little to no programming knowledge and only a basic understanding of MALDI mass spectra data analysis. We truly believe that its flexibility makes it a powerful tool for ZooMS research. The goal is to free the researcher’s mind and allow them to concentrate on more fundamental tasks.

## 6. Data availability

PAMPA: The software is open-source (license GPL3). The program together with the database of peptide markers, the COL1A1, COL1A2 sequences and the NCBI taxonomy are available from https://github.com/touzet/pampa.

All mass spectra, Fasta databases used in the case studies, csv files created for treatment with MASCOT and PAMPA and MASCOT results have been deposited on the ProteomeXchange Consortium (http://proteomecentral.proteomexchange.org) via the PRIDE partner repository with the data set identifier PXD050532 (Private data, Username, Password: 1EVJzsqS).

## Supporting information

Supp. Data: Result files

Supp. Data: description

## 7. Supplementary material

The archive supplementary_material.zip contains a description of the supplementary material (Supplementary_material _S1.pdf) as well as all result files generated by PAMPA.

## AUTHOR INFORMATION

### Author Contributions

F.B and HT: Conceptualization, writing-original draft, project administration, funding acquisition. F.B: proteomics data generation. F.B and H.T: proteomics data analyses in case studies. H.T: methodology, algorithms and software development. Both authors have given approval to the final version of the manuscript

## ACKNOWLEDGMENT

The authors warmly thank Dr Patrick Auguste from UMR 8198 – EEP - Evolution, Ecology and Paleontology, Dr Tarek Oueslati from UMR 8164 – HALMA - Histoire, Archéologie et Littérature des Mondes Anciens for providing the samples for analysis, as well as Dr Céline Poux and Dr Julie Jacques for fruitful discussions.

