## Supplementary material for "PAMPA: a software for peptide markers and taxonomic identification for ZooMS samples in Archaeology and Paleontology": Supp. Data: description

The mammal peptide table (file table_mammals.tsv), the mammal taxonomy (file taxonomy_mammals.tsv) as well as the set of COL1A1 and COL1A2 NCBI sequences are available on https://github.com/touzet/pampa
The mammal taxonomy was downloaded from Uniprot.

### 3.1 Classification of MALDI-TOF and MALDI FT-ICR modern spectra – folder Moderns

#### 3.1.1 Spectral files

The MALDI-TOF and MALDI FTICR spectra have been deposited on the ProteomeXchange Consortium (http://proteomecentral.proteomexchange.org) via the PRIDE partner repository with the data set identifier PXD050532.

| Common name | Scientific name | TOF spectra | FTICR spectra | Element & Portion | Accession number | Source |
|---|---|---|---|---|---|---|
| Bison | *Bison bison* | Bison-TOF.csv | Bison-FT.csv | Tooth, incisive | BiT1 | MSAP |
| Horse | *Equus caballus* | Horse-TOF.csv | Horse-FT.csv | Skull | HoS1 | EEP |
| Goat | *Capra hircus* | Goat-TOF.csv | Goat-FT.csv | Mandible | GoM1 | EEP |
| Cattle | *Bos taurus* | Ox-TOF.csv | Ox-FT.csv | Mandible | OxM1 | EEP |
| Sheep | *Ovis aries* | Sheep-TOF.csv | Ovis-FT.csv | Mandible | ShM1 | EEP |
| Dog | *Canis lupus familiaris* | Dog-TOF.csv | Dog-FT.csv | Skull | DoS1 | EEP |
| Fox | *Vulpes vulpes* | Vulpes-TOF.csv | Vulpes-FT.csv | Skull | FOS1 | EEP |
| Castor | *Castor canadensis* | Castor-TOF.csv | Castor-FT.csv | Skull | Cast1 | EEP |
| Red deer | *Cervus elaphus* | Red deer-TOF.csv | Red deer-FT.csv | Skull | DeS1 | EEP |
| Fallow deer | *Dama dama* | Fallow deer-TOF.csv | Fallow deer-FT.csv | Long bone | FaDeL1 | EEP |
| Whale | *Balaenoptera (genus)* | Whale-TOF.csv | whale-FT.csv | Mandible | WhaM1 | EEP |
| Vole | *Microtus arvalis* | Vole-TOF.csv | Vole-FT.csv | Bone | MolB1 | MSAP |
| Hedgehog | *Erinaceus europaeus* | Hedgehog-TOF.csv | Hedgehog-FT.csv | Phalange | HeP1 | EEP |
| Human | *Homo sapiens* | Homo-TOF.csv | Homo-FT.csv | Skull | HuS1 | EEP |
| Rabbit | *Oryctolagus cuniculus* | Rabbit-TOF.csv | Rabbit-FT.csv | Bone | RaL1 | EEP |
| Mouse | *Mus musculus* | Mouse-TOF.csv | Mouse-FT.csv | Bone | MouB1 | EEP |
| Rat | *Rattus rattus* | Rattus-TOF.csv | Rattus-FT.csv | Bone | RaB1 | EEP |
| Pig | *Sus scrofa* | Sus-dom-TOF.csv | Sus-dom-FT.csv | Mandible | PoM1 | HALMA |
| Mole | *Talpa (genus)* | Mole-TOF.csv | Moleo.d.csv | Bone | MolB1 | MSAP |

EEP : UMR 8198, Evolution, Ecology and Paleontology), University of Lille

HALMA : UMR 8164, Histoire, Archéologie et Littérature des Mondes Anciens, University of Lille

MSAP :UAR 3290 - Miniaturisation pour la Synthèse, l'Analyse et la Protéomique, University of Lille

#### 3.1.2 Analysis in marker mode

**Peptide table** : table_mammals.tsv

**Taxonomy** : taxonomy_mammals.tsv

**Command line for MALDI-TOF spectra**

```
python3 pampa_classify.py -s Spectra/TOF -p table_mammals.tsv -e 0.1
-o pampa_moderns_TOF.tsv -t taxonomy_mammals.tsv
```

**Command line for MALDI-FTICR spectra**

```
python3 pampa_classify.py -s Spectra/FT -p table_mammals.tsv -e 0.01
-o pampa_moderns_FT.tsv -t taxonomy_mammals.tsv
```

The difference between the two command-lines is the -e parameter (mass error tolerance, in Daltons here), which is lowest for MALDI-FTICR spectra.

#### 3.1.3 Analysis in all peptide mode

**NCBI sequences :** available on https://github.com/touzet/pampa_sequences/

**Taxonomy** : taxonomy_mammals.tsv

**Command line for MALDI-TOF spectra**

```
python3 pampa_classify.py -s Spectra/TOF -d ../Sequences -e 0.1
-o pampa_moderns_all_TOF.tsv -t taxonomy_mammals.tsv
```

**Command line for MALDI-FTICR spectra**

```
pampa_classify.py -s Spectra/FT -d ../Sequences -e 0.01
-o pampa_moderns_all_FT.tsv -t taxonomy_mammals.tsv
```

#### 3.1.4 Results

PAMPA result files are in the archive:

- Marker mode, TOF spectra: pampa_moderns_TOF.tsv
- Marker mode, FTICR spectra : pampa_moderns_FT.tsv
- All peptide mode, TOF spectra : pampa_moderns_all_TOF.tsv
- All peptide mode, FTICR spectra : pampa_moderns_all_FT.tsv

### 3.2 Analysis of 104 archaeological MALDI FTICR mass spectra - folder Caours

#### 3.2.1 Spectral files

This case study comes from [Bray2023]. Raw MALDI-FTICR files are available from the PRIDE database with accession number PXD038283. Processed spectra are on also on PRIDE PRIDE with accession number PXD050532.

#### 3.2.2 PAMPA command line

```
python3 pampa_classify.py -p table_mammals.tsv -s Spectra -e 0.01 --deamidation
-l deamidation_caours.txt -o pampa_caours.tsv
```

where the deamidation_caours.txt file is a single-line file that restricts application of deamidation to COL1A2-978/A and COL1A1-508/P1 markers :

```
Deamidation= COL1A2-978/A, COL1A1-508/P1
```

#### 3.2.3 Results

The result file generated by PAMPA is pampa_caours.tsv. Additionnaly, the archive contains a manual annotation of the results, spectrum by spectrum : annotation_caours.tsv. In this file, the "PAMPA" column recalls the classification assigned by PAMPA, while "Reference" corresponds to the classification reported in [Bray2023] (file Metadata_Caours.xlsx from the Supplementary Material). The "Source" column indicates whether the initial classification was based on morphology or determined by ZooMS.

### 3.3 Taxonomic identification of sheep and goat datasets – folder Sheeps_and_Goats

#### 3.3.1 Spectral files

All raw MS triplicate spectra were downloaded from https://zenodo.org/records/6967158. We selected samples having exactly three replicates: UoC01 to UoC33.

**Preprocessing for consensus spectra** : we used the R script preprocessing.md from the github repository associated to [Viñas-Caron2023] (https://github.com/ismaRP/a_spanish_book/)

**Preprocessing for individual spectra** : raw spectra also processed individually with mMass.
All resulting files are on PRIDE (33 consensus files and 99 individual files) with accession number PXD050532.

#### 3.3.2 PAMPA command-lines

Marker mode, individual analysis
```
python3 pampa_classify.py -s ../Spectra/Individual -l limit_goat_sheep.txt --
deamidation -e 0.1 -p table_mammals.tsv -o pampa_sheep_and_goat_individual.tsv
```

All peptide mode, individual analysis
```
python3 pampa_classify.py -s ../Spectra/Individual -l limit_goat_sheep.txt --
deamidation -e 0.1 -d Sequences
-o pampa_sheep_and_goat_allpeptides_individual.tsv
```

Marker mode, consensus analysis
```
python3 pampa_classify.py -s ../Spectra/Consensus -l limit_goat_sheep.txt --
deamidation -e 0.1 -p table_mammals.tsv -o pampa_sheep_and_goat_consensus.tsv
```

All peptide mode, consensus analysis
```
python3 pampa_classify.py -s ../Spectra/Consensus -l limit_goat_sheep.txt --
deamidation -e 0.1 -d Sequences
-o pampa_sheep_and_goat_allpeptides_consensus.tsv
```

#### 3.3.3 PAMPA results

- consensus spectra, marker mode : `pampa_sheep_and_goat_consensus.tsv`
- consensus spectra, all peptide mode : `pampa_sheep_and_goat_allpeptides_consensus.tsv`
- individual analysis, marker mode : `pampa_sheep_and_goat_individual.tsv`
- individual analysis, all peptide mode : `pampa_sheep_and_goat_allpeptides_invidual.tsv`

The annotation_sheeps_and_goats.tsv file provides a synthesis for all four results, with comparison to the reference classification (taken from Supplementary Data S1 of [Viñas-Caron2023] ).

### 3.4 Taxonomic identification of African bovids – folder African_Bovids

#### 3.4.1 Spectral files

99 MALDI-TOF spectra taken from https://zenodo.org/records/3971142 (three missing compared to african_bovids_supp_publi: KAV100, KAV442, KAV445) and successfully processed by SpecieScan (step 1 and step 2). Files are deposited on PRIDE, accession number PXD050532.

#### 3.4.2 Peptide table

We created a custom table, specifically designed for this example to use the same list of species and peptide markers as SpecieScan (available at https://github.com/mesve/SpecieScan/blob/main/Reference_databases/Mammals_reference_Africa.csv)

List of species :

Aepyceros melampus
Alcelaphus buselaphus
Ammotragus lervia
Antidorcas marsupialis
Bos taurus
Bubalus bubalis
Canis lupus familiaris
Capra hircus
Ceratotherium simum
Chlorocebus sabaeus
Connochaetes taurinus
Damaliscus lunatus
Diceros bicornis
Equus asinus
Equus caballus
*Eudorcas thomsonii*
*Felis catus*
*Giraffa giraffa*
*Gorilla gorilla*
*Hexaprotodon liberiensis*
*Hippopotamus amphibius*
*Hippotragus equinus*
*Hippotragus niger*
*Homo sapiens*
*Kobus ellipsiprymnus*
*Kobus kob*
*Lepus*
*Leucocephalophus adersi*
*Loxodonta africana*
*Madoqua kirkii*
*Mus musculus*
*Nanger granti*
*Neotragus moschatus*
*Oreotragus oreotragus*
*Oryctolagus cuniculus*
*Ourebia ourebi*
*Ovis aries*
*Pan paniscus*
*Pan troglodytes*
*Papio hamadryas*
*Pelea capreolus*
*Philantomba monticola*
*Pudu*
*Raphicerus campestris*
*Rattus norvegicus*
*Rattus rattus*
*Rhinolophus ferrumequinum*
*Rhinolophus hipposideros*
*Rupicapra rupicapra*
*Sus scrofa*
*Sylvicapra grimmia*
*Syncerus caffer*
*Taurotragus derbianus*
*Tragelaphus buxtoni*
*Tragelaphus eurycerus*
*Tragelaphus scriptus*
*Tragelaphus spekii*
*Tragelaphus strepsiceros*

The petide table is in the file `table_africanbovids.tsv`. Masses were taken from the Google Sheet ZooMS markers – Published data: https://docs.google.com/spreadsheets/d/15VFw1r1ePW1U3G3bek0hm0cVFefLRi2pKVLD17QIV8

#### 3.4.3 PAMPA command-line

```
python3 pampa_classify.py  -p table_africanbovids.tsv -e 0.1 --deamidation -o results_africanbovids.tsv
```

#### 3.4.4 Results

Pampa result file is `pampa_africanbovids.tsv`
These results were compared to

- the initial classification of [Janzen2021] available in the supporting information section, taken from https://doi.org/10.1371/journal.pone.0251061.s005.xlsx )
- SpecieScan results, such as made available in the file Zambia_Speciescan_results.csv taken from https://github.com/mesve/SpecieScan/blob/main/Paper%20Results%20csvs/Zambia_Speciescan_results.csv

The resulting synthesis, spectrum by spectrum, is available in the file `annotation_africanbovids.tsv`

### 3.5 Detection of mixtures – folder Mixtures

#### 3.5.1 Spectral files

Files are available on PRIDE, accession number PXD050532.

| | |
|---|---|
| bos-capra_0_C5_000001.d.csv<br>bos-capra_0_C5_000002.d.csv | *Bos taurus + Capra hircus* |
| capra-ovis_0_C7_000001.d.csv<br>capra-ovis_0_C7_000002.d.csv | *Capra hircus + Ovis aries* |
| ovis-cervus_0_M5_000001.d.csv<br>ovis-cervus_0_O5_000001.d.csv | *Ovis aries + Cervus elaphus* |

#### 3.5.2 PAMPA command-line

```
python3 pampa_classify.py -n 85 -s Spectra -e 0.01 -p table_mammals.tsv
-o pampa_mixture.tsv
```

The parameter -n 85 specifies that all suboptimal solutions with at least 85% of the matching markers found in the optimal solution are selected.

#### 3.5.3 PAMPA result file

`pampa_mixture.tsv`

### 3.6 Identification of new peptide markers by homology – folder Marine_mammals

#### 3.6.1 Peptide markers used as reference for homology

##### 3.6.1.1 For pinnipeds

table_reference_pinnipeds

| Taxon name | Sequence | Marker | PTM | Mass | Gene |
|---|---|---|---|---|---|
| Odobenus rosmarus divergens | TGHPGTVGPAGVR | A | 1H | 1221.63346016001 | COL1A2 |
| Odobenus rosmarus divergens | GLPGEFGLPGPAGPR | B | 2H | 1453.74340491225 | COL1A2 |
| Odobenus rosmarus divergens | GPPGESGAAGPSGPIGSR | C | 1H | 1566.75067474022 | COL1A2 |
| Odobenus rosmarus divergens | GLPGVSGSVGEPGPLGIAGPSGAR | D | 2H | 2121.0934729616 | COL1A2 |
| Odobenus rosmarus divergens | GLTGPIGPPGPAGAPGDKGEAGPSGPAGPTGAR | F | 2H | 2853.41257523584 | COL1A1 |
| Odobenus rosmarus divergens | GPSGEPGTAGPPGTTGPQGLLGSPGILGLPGSR | G | 4H | 3003.5017849239402 | COL1A2 |
| Odobenus rosmarus divergens | GETGPAGRPGEVGPPGPPGPTGEK | P | 3H | 2246.06838079301 | COL1A1 |
| Odobenus rosmarus divergens | GVQGPPGPAGPR | Cet1/P1 | 1H | 1105.57488265473 | COL1A1 |

File in the archive : table_reference_pinnipeds.tsv

##### 3.6.1.2 For cetaceans

table_reference_cetaceans

| Taxon name | Sequence | PTM | Mass | Marker | Gene |
|---|---|---|---|---|---|
| Balaenoptera musculus | TGHPGAVGPAGIR | 1H | 1205.63854554045 | A | COL1A2 |
| Balaenoptera musculus | GIPGEFGLPGPAGPR | 2H | 1453.74340491225 | B | COL1A2 |
| Balaenoptera musculus | GPPGESGAAGPAGPIGNR | 1H | 1577.76665915753 | C | COL1A2 |
| Balaenoptera musculus | GLPGVAGSVGEPGPLGISGPAGAR | 2H | 2105.0985583420397 | D | COL1A2 |
| Balaenoptera musculus | GLTGPIGPPGPAGAPGDKGETGPSGPAGPTGAR | 2H | 2883.4231399195396 | F | COL1A1 |
| Balaenoptera musculus | GPSGEPGTAGSPGTPGPQGLLGAPGFLGLPGSR | 5H | 3023.4704851760994 | G | COL1A2 |
| Balaenoptera musculus | GTTGEIGSAGPPGPPGLR | 2H | 1652.82384006092 | Cet2/P2 | COL1A2 |
| Balaenoptera musculus | GVQGPSGPAGPR | 0H | 1079.55923221015 | Cet1/P1 | COL1A1 |
| Tursiops truncatus | TGHPGAVGPAGIR | 1H | 1205.63854554045 | A | COL1A2 |
| Tursiops truncatus | GIPGEFGLPGPAGPR | 2H | 1453.74340491225 | B | COL1A2 |
| Tursiops truncatus | GPPGESGAAGPTGPVGSR | 1H | 1566.75067474022 | C | COL1A2 |
| Tursiops truncatus | GLPGVAGSVGEPGPLGIAGPTGAR | 2H | 2119.1142084061803 | D | COL1A2 |
| Tursiops truncatus | GLTGPIGPPGPAGAPGDKGETGPSGPAGPTGAR | 2H | 2883.42313991954 | F | COL1A1 |
| Tursiops truncatus | GPSGEPGTAGSPGTPGPQGLLGAPGFLGLPGSR | 5H | 3023.4704851760994 | G | COL1A2 |
| Tursiops truncatus | GSTGEIGSAGPPGPPGLR | 2H | 1638.80818999678 | Cet2/P2 | COL1A2 |
| Tursiops truncatus | GVQGPSGPAGPR | 0H | 1079.55923221015 | Cet1/P1 | COL1A1 |

File in the archive : table_reference_cetaceans.tsv

#### 3.6.2 NCBI sequences

| | Family | *Species or subspecies* | COL1A1 | COL1A2 |
|---|---|---|---|---|
| Mysticeti | Balaenopteridae | *Balaenoptera ricei* | XP_059763561.1 | XP_059789839.1 |
| Mysticeti | Balaenidae | *Eubalaena glacialis* | XP_061030802.1 | XP_061030802.1 |
| Mysticeti | Balaenopteridae | *Balaenoptera acutorostrata* | XP_057391191.1 | XP_007195818.2 |
| Mysticeti | Balaenopteridae | *Balaenoptera musculus* | XP_036691630.1 | XP_036718747.1 |
| Odontoceti | Delphinidae | *Tursiops truncatus* | XP_033703059.1 | XP_019783093.1 |
| Odontoceti | Kogiidae | *Kogia breviceps* | XP_058902606.1 | XP_058929757.1 |
| Odontoceti | Phocoenidae | *Neophocaena asiaeorientalis asiaeorientalis* | XP_024601210.1 | XP_024622854.1 |
| Odontoceti | Phocoenidae | *Phocoena sinus* | XP_032471594.1 | XP_032498638.1 |
| Odontoceti | Physeteridae | *Physeter catodon* | XP_007128607.1 | XP_007128607.1 |
| Odontoceti | Ziphiidae | *Mesoplodon densirostris* | XP_059938244.1 | XP_059963578.1 |
| Odontoceti | Delphinidae | *Delphinus delphis* | XP_059853534.1 | XP_059876500.1 |
| Odontoceti | Delphinidae | *Globicephala melas* | XP_030695607.1 | XP_030713115.2 |
| Odontoceti | Delphinidae | *Lagenorhynchus obliquidens* | XP_026950147.1 | XP_026958275.1 |
| Odontoceti | Delphinidae | *Orcinus orca* | XP_004282670.1 | XP_004265635.1 |
| Odontoceti | Monodontidae | *Delphinapterus leucas* | XP_022413783.1 | XP_022441863.1 |
| Odontoceti | Monodontidae | *Monodon monoceros* | XP_029068332.1 | XP_029089490.1 |
| Pinnipeds | Otariidae | *Eumetopias jubatus* | XP_027949502.1 | XP_027974643.1 |
| Pinnipeds | Otariidae | *Zalophus californianus* | XP_027423360.2 | XP_027428797.1 |
| Pinnipeds | Phocidae | *Mirounga angustirostris* | XP_045745072.1 | XP_045720640.1 |
| Pinnipeds | Odobenidae | *Odobenus rosmarus divergens* | XP_004395207.1 | XP_004394135.1 |

| | | | | |
|---|---|---|---|---|
| Pinnipeds | Otariidae | *Callorhinus ursinus* | XP_025715155.1 | XP_025715155.1 |
| Pinnipeds | Phocidae | *Halichoerus grypus* | XP_035950567.1 | XP_035979250.1 |
| Pinnipeds | Phocidae | *Mirounga leonina* | XP_034882768.1 | XP_034876481.1 |
| Pinnipeds | Phocidae | *Phoca vitulina* | XP_032284949.1 | XP_032258023.1 |
| Pinnipeds | Phocidae | *Neomonachus schauinslandi* | XP_021560003.1 | XP_021545228.1 |

The sequences are available on https://github.com/touzet/pampa_sequences

#### 3.6.3 PAMPA command lines

**For pinnipeds**

```
python3 pampa_craft.py --homology -p table_reference_pinnipeds.tsv -d  Sequences
-l limit_pinnnipeds.txt -o table_homology_pinnipeds.tsv -t
../Taxonomy/taxonomy_mammals.tsv
```

where the file limit_pinnipeds.txt is the single-line file

```
OX=9705, 9702, 9709
```

**For cetaceans**

```
python3 pampa_craft.py --homology -p table_reference_cetaceans.tsv -d  Sequences
-l limit_cetaceans.txt -o table_homology_cetaceans.tsv -t
../Taxonomy/taxonomy_mammals.tsv
```

where the file `limit_cetaceans.txt` is the single-line file

```
OS=Cetacea
```

#### 3.6.4 Result files

The two files  pampa_homology_pinnipeds.tsv and pampa_homology_cetaceans.tsv are available in the archive.

### 3.7 De novo full peptide generation and filtering  - folder Sheep_Filtering

#### 3.7.1 Analysis with all tryptic peptides

Command-line

```
python3 pampa_craft.py --allpeptides -f sheep.fasta -o
pampa_sheep_allpeptides.tsv
```

where sheep.fasta contains the COL1A1 and COL1A2 sequences for *Ovis aries*.

Result file :  `pampa_sheep_allpeptide.tsv`

#### 3.7.2 Analysis with peak filtering

Spectra : 57 files individual maldi-TOF spectra coming from the 19 samples from  [Viñas-Caron2023] annotated as sheep (UcCO1, UcCO2, UcCO3, UcCO4, UcCO6, UcCO13, UcCO14, UcCO15, UcCO15, UcCO17, UcCO17, UcCO18, UcCO19,UcCO20, UcCO21, UcCO22, UcCO23, UcCO24, UcCO25 )

Command-line

```
python3 pampa_craft.py --allpeptides -f sheep.fasta -o pampa_sheep_filter.tsv -s
Spectra -e 0.1
```

Result file : `pampa_sheep_filtering.tsv`
